# ZeaMiC: a Publicly Available Culture Collection of Maize Root-Associated Bacteria

**DOI:** 10.64898/2026.03.23.713778

**Authors:** Anna-Katharina Garrell, Nichole Ginnan, Joel F. Swift, Gaurav Pal, Athanasios Zervas, Christine Pestalozzi, Clara Tang, Felicity Tso, Natalie E. Ford, Ben Niu, Gabriel Castrillo, Klaus Schläppi, Richard L. Hahnke, Maggie R. Wagner, Manuel Kleiner

## Abstract

Plant-associated microbiota are composed of hundreds of microbial species. For many of them, little is known about their individual functions and even less is known about their emergent community-level traits. While culture-independent methods provide valuable insights into the composition, diversity, and functional potential of plant-associated microbiota, culture-dependent methods are essential for reductionist lines of inquiry into the roles of individual species and their interactions within a community. Here, we present ZeaMiC, a publicly available culture collection of root-associated bacteria from *Zea mays* (maize). This resource comprises 88 isolates obtained from diverse soils and several maize genotypes, with live cultures available through DSMZ (German Collection of Microorganisms and Cell Cultures) both as single stocks and as cost-effective bundles (https://www.dsmz.de/collection/catalogue/microorganisms/microbiota/zeamic). To maximize relevance, isolates were selected to be representative of maize root-associated microbiomes in the Corn Belt of the United States, based on abundance-occupancy patterns from previously published root microbiome data, phylogenetic diversity, and literature-based evidence of functional importance. Whole-genome sequencing and annotation revealed genes associated with root colonization, plant growth promotion, and nutrient cycling, including functions such as chemotaxis, biofilm formation, secretion systems, hormone modulation, and phosphate solubilization. This collection serves as a community resource for future mechanistic studies of plant-microbe and microbe-microbe interactions, filling the gap in our understanding of the ecological interactions in plant microbiomes.

## Introduction

Plant-associated microbial communities are increasingly recognized as integral components of agricultural systems, influencing plant health, productivity, and resilience to stress ^1–4^. For staple crops like maize (*Zea mays*) or corn, which ranks among the top globally produced crops ^5,6^, deciphering the structure and function of root-associated microbial communities has direct implications for global food security, animal feed, and bioenergy production. Advances in high-throughput sequencing have enabled detailed surveys of maize root microbiota, providing insight into their taxonomic composition across diverse environments. These studies have revealed that maize root-associated microbial communities have consistent taxonomic profiles ^7–18^.

However, despite advances in culture-independent techniques, much of our understanding of the maize rhizosphere microbiome remains descriptive, lacking direct experimental investigation of plant-microbe interactions. Axenic cultures of host-associated microbes are indispensable for studying host-microbe and microbe-microbe interactions under controlled conditions, allowing for the direct testing of mechanistic and ecological hypotheses. Microbial culture collections thus complement culture-independent datasets by providing a foundation for rational testing of hypotheses, both in single species- and in more complex, multi-species systems ^19–21^. For example, the *Arabidopsis* phyllosphere and rhizosphere culture collections (*At*-LSPHERE and *At*-RSPHERE) ^22^ have enabled studies of mechanisms driving microbe-microbe interactions *in planta*, microbiome-mediated protection against plant pathogens, host response to bacterial inoculation, and community assembly ^23–25^. Similar culture collections have also been developed for a variety of crops, including barley ^26^, sugarcane ^27^, and switchgrass ^28^, among others ^25,26,29–32^.

In maize, a collection of root-associated bacteria developed by Thoenen *et al.* ^33^ has supported the study of microbial metabolism of host-derived compounds ^34,35^ and a synthetic community of root-associated bacteria developed by Niu *et al.* has been used to investigate community structure ^9^, microbe-dependent heterosis ^36^, and bacterial functions activated when grown in root-associated environments ^37^. Additionally, Custódio *et al.* developed a collection of maize root- and leaf-associated bacteria, which were used to investigate the effects of bacterial inoculation on leaf growth and host transcriptional response ^38^.

Despite these successful efforts to collect and identify maize-associated microbial isolates, the availability of cultures originating from maize grown in the Midwest United States (US) is noticeably lacking. The US is the global leader of corn production, producing 31% of the world’s corn ^39^. Thirteen US states, including Kansas and Michigan, account for approximately 90% of the national corn production ^40^. The majority of these states are in the Midwest region and are commonly referred to as the “Corn Belt”, which is considered the world’s most productive maize-producing region. Thus, the absence of publicly available isolates from this region potentially limits our ability to explore how maize interacts with some of its most frequently encountered microbial neighbors.

In an effort to create a comprehensive and publicly available collection of maize root-associated bacteria reflective of US Corn Belt microbiomes, we have created a taxonomically and functionally diverse culture collection of 88 maize root-associated bacteria, ZeaMiC (**Zea** mays **Mi**crobial **C**ollection). We designed this collection to represent the core microbiota previously described for maize grown in prairie and agricultural soils from the US Corn Belt ^41^, and supplemented it with previously described isolates from maize grown in Swiss agricultural soils ^33^, US horticultural soils ^9^, and United Kingdom and Cape Verde agricultural soils ^38^. We further show that the bacterial isolates in this collection are commonly found across numerous maize microbiome studies from across the globe, with a substantial proportion of the relative abundance sampled within these studies captured by ZeaMiC isolates. ZeaMiC currently includes only bacteria, though we envision the resource growing in the future to include fungi and other microbes. We have made this collection available through the German Collection of Microorganisms and Cell Cultures (DSMZ) both as individual strains and cost-effective isolate bundles, and the associated genomes are available through the National Center for Biotechnology Information (NCBI). This collection represents an international, multi-laboratory effort to develop a publicly available community research tool that covers most of the diversity of bacterial taxa associated with maize roots.

## Materials and Methods

### Overview: Collection rationale and design

Our goal was to generate a community-accessible, experimentally tractable bacterial culture collection that is representative of the maize root bacteriome and suitable for mechanistic studies of the maize microbiome (Fig. 1).

**Figure 1.**
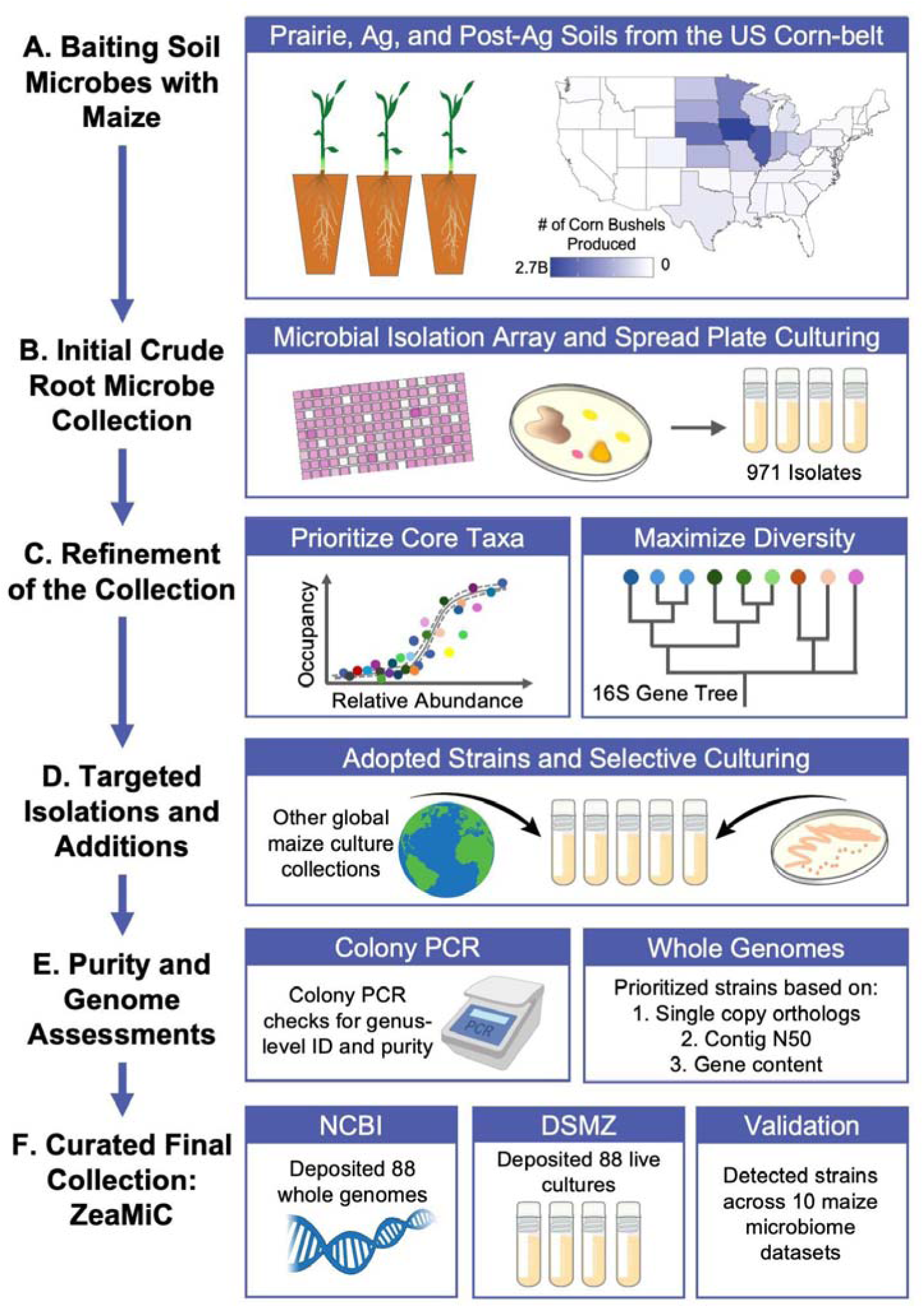
Workflow for the construction of a maize root bacterial culture collection (ZeaMiC). **(A)** Soil microbiota from prairie, agricultural, and post-agricultural fields were used to inoculate maize seedlings and “bait” root colonizing bacteria. Corn bushel production map was produced using USDA 2025 data (https://quickstats.nass.usda.gov/). **(B)** Root-colonizing bacteria were isolated using high-throughput array and spread plating approaches, yielding >900 isolates that were purified and identified at the genus-level by full-length 16S rRNA gene sequencing. **(C)** Core taxa were prioritized based on abundance–occupancy patterns from a prior maize root microbiome dataset, and a 16S rRNA gene phylogeny of 971 isolates was used to maximize phylogenetic diversity in the refined collection. **(D)** Targeted isolation and incorporation of strains from existing maize culture collections increased recovery of priority taxa not captured initially. **(E)** Selected strains underwent purity and taxonomic validation followed by whole-genome sequencing, and genome quality metrics guided final selection. **(F)** The resulting 88-strain collection was deposited in public repositories (NCBI GenBank and DSMZ), and strain relevance was validated across previously published maize microbiome datasets.

Using a previously published *16S rRNA* gene amplicon dataset ^41^ from the roots of maize plants grown in microbial communities originating from US Midwest soils, we defined core bacterial genera as those exhibiting both high relative abundance and high prevalence across maize root samples. This abundance-occupancy approach has been shown to identify persistent host-associated taxa and provides a practical strategy for prioritizing bacteria most likely to contribute to reproducible colonization and community function ^42^. Furthermore, studies in animal gut-associated microbiomes indicate that core taxa often possess larger genomes and greater metabolic independence than non-core members, suggesting that communities enriched for core taxa can capture key functional attributes of more complex microbiomes ^43^.

Initial bacterial isolates were derived from soils collected in the US Midwest region (Fig. 1A). Maize seedlings were grown with these soil microbiota and then root-colonizing bacteria were isolated using both high-throughput microbial culture arrays and standard spread plating approaches (Fig. 1B). While selecting strains for the final culture collection we aimed to (i) maximize the overall taxonomic diversity of the collection and (ii) target core bacterial taxa identified in the abundance-occupancy analysis (Fig. 1C). To improve taxonomic coverage in both of these areas, we also incorporated previously published bacterial isolates from maize roots grown in soils from geographic regions across the world (Fig. 1D). Together, these efforts produced the ZeaMiC, a refined, genomically characterized, centralized maize root bacterial culture collection that is available to the research community through the DSMZ (Fig. 1F). We validated the relevance of ZeaMiC bacterial isolates by quantifying their presence and relative abundance across ten published maize microbiome global datasets.

### Generation of crude isolate collection

#### Soil inoculum collection

Soil samples were collected from six field sites in the US Midwest that represent prairie, post-agricultural, and agricultural microbial communities. These include The Land Institute (TLI), Konza Wildlife Reserve (KNZ), Hays Prairie (HAY), and Smoky Valley Ranch (SVR), the University of Kansas Field Station (KU), and the Michigan State University (MSU) plant pathology research farm. Soil sample site metadata including sampling dates, locations, and management information are available in Supplementary Table S1.

TLI, KNZ, HAY, SVR, and KU fields were split into four subplots. In each subplot, the top 10 cm of surface soil and thick plant root masses were removed with a sterilized metal shovel. Then, ∼1 L of soil was collected from each subplot, pooled, and homogenized in a polyethylene bag, for a total of ∼4 L of soil collected per site. Soil was held at room temperature during transport back to the laboratory where it was then stored at 4°C. For the MSU soil, ∼200 g of soil was collected from four random locations within two separate agricultural fields (targeting the spaces between plants) and then mixed together into one final sample. The MSU soil was collected to a depth of ∼5 cm.

#### Inoculating maize with field soil

Soil slurries were created by placing 20 g of soil in 100 mL of 1x phosphate-buffered saline (PBS) with 0.0001% Triton X-100 and shaken at 300 rpm at room temperature for 10 mins. Soil solutions were filtered through sterile miracloth (Millipore Sigma, #475855, Burlington, MA, USA) into 50 mL conical tubes. Tubes were centrifuged for 30 mins at 3000 xg. The supernatant was removed and the pellet was resuspended in 20 mL of 1x PBS. The soil slurry was diluted further by adding 5 mL of slurry to 500 mL of 0.5x Murashige-Skoog (MS) liquid media, resulting in 0.01 g of soil per mL after dilution.

Maize seeds (genotype B73) were surface-sterilized by submerging in 70% ethanol for three mins, then 5% sodium hypochlorite for two mins, and finally rinsed in sterile water three times. Surface-sterilized seeds were planted in 100 mL cone-tainers (Stuewe & Sons, #SC10R, Tangent, OR) that contained either sterile calcined clay (Oil Dri, #44Z111, Chicago, IL) inoculated with 25 mL of a field soil slurry, or a 4:4:1 mixture of autoclaved sand, autoclaved vermiculite, and field soil. For the MSU and KU soils, we only used the 4:4:1 mixture method because Gammaproteobacteria were being isolated at a higher rate than expected from the soil slurry-inoculated plants, and it is likely that the slurry method was enriching specific microbial taxa. Plants were placed in a growth chamber set to a 12-hr day cycle, 27°C/23°C, and ambient humidity for one month. We watered all plants two times per week to maintain well watered conditions. No fertilizer was applied.

#### Isolation of maize root bacteria

One-month-old maize plants were harvested from pots and soil/clay adhered to the root surface was gently shaken off. To enrich for root endophytes (microbes that inhabit internal root niches), we surface-sterilized roots by submerging in 95% ethanol and gently shaking for 1 min, then submerging in 3% sodium hypochlorite for 5 mins, then submerging in 95% ethanol for 1 min, and finally rinsing with sterile water three times. We isolated bacteria from the surface-sterilized roots using two main approaches, (i) high-throughput microbial cultivation array technology and (ii) spread plating on Petri dishes.

For high-throughput microbial isolations, three replicates (whole root systems) from the same soil source were pooled and then cut into ≤ 1 cm fragments using a flame sterilized razor blade. Between 2-2.5 g of root fragments for each sample were placed into separate 15 mL conical tubes. Tubes were then filled with sterile 25% glycerol, incubated at room temperature for 20 mins, and then placed at -20°C until they were shipped to General Automation Lab Technologies (GALT; now Isolation Bio, Inc., San Carlos, CA, USA) for isolation in 50% strength R2A media using their Prospector microbial isolation array technology. Bacterial cultures were received from GALT as glycerol stocks in multiple 96-well plates, with each well presumably containing a single bacterial strain.

For spread plate isolations using Petri dishes, three surface-sterilized replicates (whole root systems) were placed into a mesh extraction bag (Agdia, ACC 00930/0100, Elkhart, IN, USA) containing 1 mL of 1x PBS buffer and crushed using a hammer. The resulting root slurries were diluted 1:10, 1:100, 1:1000, and 1:10000 with 1x PBS buffer and 100 µL of each dilution was spread onto 1x R2A agar plates (Fisher Scientific, NEOGEN 700004593). Plates were incubated at 28°C for 5 days. Bacterial strains from maize plants grown in MSU and KU 4:4:1 (sand:vermiculite:field soil) mixtures were all isolated using the spread plate approach (Supplementary Table S2).

#### Identification and storage of root bacterial isolates

The glycerol stocks generated by GALT, which were expected to have one strain per well, and colonies from the spread plates, with different morphologies, were streaked onto separate 1x R2A medium plates and subcultured until pure isolated colonies were obtained (Supplementary Table S2). Colony PCR was performed for each pure isolate to amplify the *16S rRNA* gene using the 27F (5’-AGAGTTTGATCCTGGCTCAG-3’) and 1492R (5’-GGTTACCTTGTTACGACTT-3’) primer set. Clean-up and Sanger sequencing of crude PCR products were performed by Genewiz (South Plainville, NJ, USA). We sequenced with only the forward primer to reduce costs of this preliminary screening step. The sequences obtained from Genewiz therefore ranged from 800-1200 bp rather than the full length of the *16S rRNA* gene (∼1500 bp); we thus refer to them as partial *16S rRNA* gene sequences. Isolated colonies were inoculated and grown in R2A broth at 28°C and 200 rpm for 24-72 hrs until the culture was turbid. Liquid cultures were used to make individual glycerol stocks at a 1:1 ratio of sterile 50% glycerol to liquid culture. These 971 isolates comprised the “crude” culture collection (Supplementary Table S2).

### Selection of high-priority candidate strains from the crude isolate collection

Our high-throughput cultivation array and agar-plate isolation efforts resulted in a total of 971 isolates with genus-level taxonomy assigned by aligning our partial *16S rRNA* gene sequences to the NCBI core_nt reference database using NCBI BLASTn ^44^ (Supplementary Table S2). We narrowed down this crude isolate collection to a smaller group of “candidate strains” that (1) maximized phylogenetic diversity of the collection and (2) prioritized representation of core taxa identified using pre-existing data on maize root microbiome composition.

#### Selecting candidate strains to maximize phylogenetic coverage

We generated a multiple sequence alignment based on the partial *16S rRNA* gene sequences for the 971 isolates in the crude collection using MUSCLE ^45^ implemented in MEGA X ^46^. The alignment was trimmed using ClipKIT’s smart-gap method ^47^ and a maximum-likelihood tree was built using a Tamura-Nei model. The tree was visualized using iTOL ^48^ (Supplementary Fig. S1). A bar plot was created to visualize the number of isolates assigned to each genus in R using ggplot2 ^49^ (Supplementary Fig. S2).

The *16S rRNA* gene tree was used to guide selection of individual strains for our de-replicated collection, ensuring that all major clades were represented in our final isolate collection (Supplementary Table S2). To start, we selected one strain per available genus, prioritizing the strain with the highest *16S rRNA* gene sequence similarity to the reference strain for the genus in NCBI. If more than one strain was available for a genus, we used the strains’ placement on the tree to select additional strains that had divergent full-length *16S rRNA* gene sequences (“candidate strains”), with the goal of capturing functional variation within genera.

#### Selecting candidate strains to represent core taxa

We used data from a previous maize *16S rRNA* gene amplicon sequencing experiment ^41^ to identify core root bacterial genera, which guided downstream selection of bacteria to include in the final culture collection. We focused on this study as it utilized the same Kansas soils that were the main sources of isolates for the collection. Briefly, maize seedlings were inoculated with soil slurries derived from Kansas prairie soils and sequencing of the 4th variable (V4) region of the *16S rRNA* gene was used to characterize root bacterial colonization of 50-day old plants.

We defined core taxa by generating an abundance-occupancy curve, selecting genera rather than species because the V4 region of the *16S rRNA* gene lacks sufficient resolution to reliably distinguish species or strains ^50^. Abundance was calculated as the average relative abundance of each genus across samples and log_10_-transformed, while occupancy was calculated as the percentage of samples (out of 200) in which the genus was present. Rather than applying strict quantitative thresholds for inclusion in the collection, we used this approach to prioritize bacterial genera with high occupancy (that is, those observed in a large proportion of plants), particularly those occurring more frequently than expected based on abundance alone ^42^. This prioritization resulted in a set of 35 core genera that together accounted for approximately 97% of the total relative abundance across samples (individual genera ranged from 0.1% to 31% relative abundance) and exhibited high occupancy (19% to 100%). We used the package fantaxtic (https://github.com/gmteunisse/Fantaxtic) to generate plots with the 35 genera.

### Targeted additions of high-priority candidate taxa

We generated a list of high-priority taxa for which we did not have isolates in our crude culture collection. Priority was based on (i) strains from taxa that were previously observed to show high abundance or occupancy in maize roots ^41^, and/or (ii) published evidence that they are associated with maize roots and/or benefits to plant growth ^9,33,38,51–53^. Bacterial taxa prioritized based on high abundance or occupancy are listed in Supplementary Table S3. We then acquired representatives of the missing high-priority taxa through (1) adoption from previously existing global maize isolate collections, or (2) targeted isolation efforts.

#### Community-enabled acquisition of isolates from high-priority taxa

To fill the taxonomic gaps in our collection, we leveraged prior isolation efforts by the global maize microbiome research community to obtain additional strains that met our criteria for inclusion. We added strains of *Methylobacterium, Arthrobacter, Paenarthrobacter, Enterobacter* (now *Franconibacter* ^54^)*, Pantoea,* and *Sinomonas* that had been previously isolated from maize roots (genotypes B73 and Zm523) grown in soils collected from the United Kingdom and Cape Verde ^38^. We also added strains of *Bacillus, Peribacillus, Deinococcus, Streptomyces, Nocardioides, Agrobacterium,* and *Sphingobium* that were derived from maize roots (genotype B73) grown in soil from an arable field in Changins, Switzerland ^33^. These strains were imported to Kansas, USA under USDA APHIS permit number 526-23-263-16301. Last, we added seven previously described strains ^9^ isolated from maize roots (genotype “Sugar Buns”) utilized in past studies of maize-microbe ^36,37^, and microbe-microbe ^55^ interactions, and for which established tools and protocols ^56,57^ are already available.

#### Targeted isolation of a Telluria sp. – a high-priority taxon

We took extra steps to isolate a *Telluria* sp., which was absent from our initial isolation effort and was not available through the global maize-microbe research community. We had identified *Telluria* as a high-priority genus based on its predominance in our maize root microbiome data (Supplementary Table S3) and its close relationship to *Massilia* ^58^, which has documented effects on important maize phenotypes ^59,60^. Isolation of *Telluria* sp. from Konza Prairie soil was accomplished following a strategy described by Niu *et al.* ^9^. Maize seeds (genotype B73) were surface-sterilized ^61^ and pre-germinated in sterile Petri plates containing 3 mL sterile water for 3 days. Germinants were transferred to four gnotobiotic bags (Whirl-Pak, Nasco Sampling, Chicago, IL, USA), three containing a 3:7 mixture of Konza Prairie soil and autoclaved calcined clay, and one containing unamended Konza Prairie soil. Each bag received 60 mL of 0.5x MS medium and was sealed with Aeraseal (Millipore Sigma) before being transferred to a growth chamber (12 hr. days, 27°C/23°C, ambient humidity). After 9 days, bags were cut open, and roots were gently shaken to remove loosely attached soil. Roots were transferred to 40 mL PBS in a 50 mL Falcon tube and vortexed for 15 seconds. The washing was repeated five times with fresh PBS until the wash remained clear. Washed roots were cut into 0.5 cm segments, pooled, and homogenized in 20 mL PBS using a sterile mortar and pestle to obtain a root slurry. This root slurry was used to inoculate another set of surface-sterilized, pre-germinated maize seedlings by incubation for 1 hour. Inoculated seedlings were transferred to glass tubes containing 0.8% MS agar, sealed with an inverted test tube and parafilm, and incubated under light emitting diode (LED) growth lights (16-hr days, 23°C, ambient humidity) for 10 days. Roots were then harvested and macerated using sterile mortar and pestle. Serial dilutions (up to 10^-5^) were prepared in triplicate. 100 µL from four dilutions (10^-2^ to 10^-5^) were plated on R2A agar (1%, 5%, and 10%) in triplicate. Plates were incubated for 3 days at 30°C. Bacterial colonies were selected based on morphology and further cultured on R2A. Isolates were identified using *16S rRNA* gene sequencing as described above. PCR products were ethanol-precipitated and sent for Sanger sequencing (Eton Bioscience, USA).

### Refinement and characterization of the final curated collection

Glycerol stocks of the selected candidate strains were shipped to the Kleiner Lab at North Carolina State University (NCSU), where the colony PCR of the *16S rRNA* gene was repeated using the GM3F (5’-AGAGTTTGATCMTGGC-3’) and GM4R (5’-TACCTTGTTACGACTT-3’) primer set to confirm identity and purity. PCR products were sent to Eton Bioscience (Research Triangle Park, NC, USA) for Sanger sequencing. Forward and reverse sequences were aligned using Clustal W^41^ and manually trimmed for quality. From there, we generated genome sequences for all candidate strains from the original US Midwest collection, which we used to de-replicate strains. Genomes for all ZeaMiC members are available through NCBI BioProjects PRJNA1250502, PRJNA1009252, or PRJNA357031 (Supplementary Table S4).

#### Whole-genome sequencing and assembly for US Midwest candidate strains

For the US Midwest (TLI, KNZ, HAY, SVR, KU, and MSU) candidate isolates, DNA extraction was performed using a Masterpure DNA complete isolation kit (Qiagen, Hilden, Germany), following the manufacturer’s protocol, except for the DNA hydration solution, where 10 mM Tris, 50 mM NaCl pH 8.0 was used instead. For the Gram+ isolates, 10 µL of lysozyme were added to the Tissue and Cell Lysis Solution. The quality of the DNA eluates was measured on a Nanodrop 2000 spectrophotometer (Thermo Scientific) and the quantity was measured on a Qubit4 fluorometer (Thermo Scientific).

Libraries for Illumina sequencing were prepared using the NexteraXT kit, following the manufacturer’s instructions. DNA was resuspended in PCR-grade water instead of re-suspension buffer. The pooled genomes were sequenced on a NextSeq 500 using the v2.5, 300-cycle chemistry in 150 bp paired-end mode (Illumina). Libraries for Oxford Nanopore Technologies (ONT) sequencing were prepared using the Rapid Barcoding kit SQK-RBK114.96, following the manufacturer’s instructions (Oxford Nanopore Technologies). Sequencing was performed on a MinION mk1b using R10.4.1 flowcells controlled by MinKNOW 22.10.10. Basecalling and trimming of barcodes and adapters were performed using dorado v0.8.4+98456f7 (https://github.com/nanoporetech/dorado).

Illumina raw reads were processed through an in-house automated pipeline (https://zenodo.org/badge/latestdoi/531835641). Briefly, raw reads were quality trimmed with TrimGalore (https://github.com/FelixKrueger/TrimGalore), then assembled with SPAdes v3.15.0 ^62^ and assembly statistics were determined with QUAST v5.2.0 ^63^. Open reading frames were predicted with PROKKA ^64^ v1.14.6. Genome comparisons were performed with dRep v3.4.0 ^65^ and checkM v1.2.1 ^66^ using the lineage_wf workflow. The assembled genomes were inspected for completion using BUSCO 5.4.0 ^67^ and its generalized bacteria_odb10 database. Full specifications of the programs used, including their versions and options are listed in the Github repository (https://github.com/AU-ENVS-Bioinformatics/IlluminaSnakemake).

ONT trimmed reads were assembled using flye v2.9-b1768 ^68^ using the --nano-hq flag, with 10 polishing iterations. The ONT assembled genomes were processed similarly to the Illumina draft genomes.

For genera containing multiple candidate strains, we used the resulting genome sequences to remove redundant strains from the final collection. Pairwise average nucleotide identity (ANI) scores were calculated for all genomes, then clustered. In cases where isolates formed a cluster with greater than 98% ANI, we examined gene content using GenAPI v1.0 and assessed completeness and assembly statistics with BUSCO v5.4.0 to select a minimal number of isolates from each cluster for inclusion in the collection. Priority was given first to isolates with complete or near-complete sets of single-copy orthologs from the bacterial lineage dataset (bacteria_odb10), second to isolates with the highest contig N50 values, and lastly, when those conditions were equivalent, to isolates based on gene content, including the number of shared and unique genes annotated by GenAPI.

#### Whole-genome sequencing and assembly for globally-sourced strains

For isolates acquired from the United Kingdom and Cape Verde ^38^, DNA extraction and whole genome sequencing were performed by Plasmidsaurus using Oxford Nanopore Technology with custom analysis and annotation.

Genomes for the *Bacillus, Peribacillus, Streptomyces, Nocardioides, Agrobacterium,* and *Sphingobium* isolates acquired from Changins, Switzerland soil were already published ^33^. *Deinococcus* LDE1, which was from the same culture collection, however, was not included in this previous sequencing effort. Therefore, DNA was extracted from *Deinococcus* LDE1 using the GenElute^TM^ Bacterial Genomic DNA Kit according to the manufacturer’s recommendations for Gram-positive bacteria. DNA concentration was measured using the AccuClear® Ultra High Sensitivity dsDNA Quantitation kit (Biotium) and DNA purity was measured using a Nanodrop One spectrophotometer (Thermo Scientific). Bacterial DNA-seq libraries were made using a plexWell 96 Library Preparation Kit for Illumina Sequencing Platforms (SeqWell, Part Nos. PW096) according to their user guide (SeqWell, v20220429) and with 10 ng of input DNA. The final library pool was evaluated using a Thermo Fisher Scientific Qubit 4.0 fluorometer with the Qubit dsDNA HS Assay Kit (Thermo Fisher Scientific, Q32854) and an Agilent Fragment Analyzer (Agilent) with a HS NGS Fragment Kit (Agilent, DNF-474), respectively. The library pool was sequenced using a NextSeq1000/2000 P1 Reagents (300 cycles, 150 bp paired-end) kit (Illumina, 20100982) on an Illumina NextSeq 1000 instrument. All steps from DNA extraction to sequencing data generation and data utility were performed at the Next Generation Sequencing Platform, University of Bern, Switzerland. Paired-end reads were quality filtered and trimmed using fastp v0.23.4 using default settings and assembled using SPAdes v4.1.0 ^62^.

#### Validation of ZeaMiC representation in global maize root microbiome studies

Ten studies were selected to assess the extent to which ZeaMiC is representative of maize microbiomes from a broad geographic distribution from across North America and Europe ^12,16–18,41,60,69–72^. While not a complete survey of the maize microbiome research corpus, these studies represented over 10,000 individual samples, hundreds of genotypes, various stressors, and multiple maize microbiome compartments. Details on sequence processing, amplicon sequence variant (ASV) filtering, and alignment of ZeaMiC taxa to the published ASVs are provided in Supplemental Appendix 1. To assess the presence and quantify the relative abundance of ZeaMiC isolates across the 10 studies, the isolates’ 16S *rRNA* sequences were aligned to the published maize-associated ASVs at 97% and 99% sequence identity. Where relevant, results were averaged across all compartments and experimental treatments within each study to provide a generalized and robust assessment; compartment-stratified values were also computed.

In addition to analyzing each study individually, we integrated ASVs across all studies to assess overall coverage of ZeaMiC and to identify candidates for future addition to the collection. We generated an abundance–occupancy curve in which ASVs were annotated based on whether they were represented in ZeaMiC at 99% sequence identity. To prioritize taxa for future inclusion, we identified genera absent from ZeaMiC that exhibited the highest mean relative abundance and prevalence across studies (present in at least 9 studies). For each study, ASV-level relative abundance and occupancy were first calculated. Then, the relative abundance of each genus (□) was calculated as the mean of study-level relative abundances of all ASVs assigned to that genus:

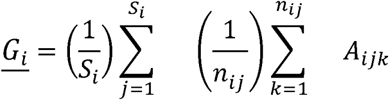

where *S_i_* is the number of studies in which genus *i* was present; *n_ij_* is the number of ASVs in genus *i* that were observed in study *j,* and *A_ijk_* is the relative abundance of the *k*^th^ ASV in genus *i* measured in study *j*. We used the abundances of genera rather than individual ASVs because sequence alignment among studies was not possible due to differences in the *16S rRNA* gene regions targeted (V3–V4, V4, and V5–V7).

#### Identification of plant-associated microbial genes

To identify microbial genes encoded by the isolates in our collection that are known to be involved in rhizosphere and root colonization ^73–77^, plant growth promotion ^74,75,78^, and nitrogen and phosphorus cycling ^76,77^, we manually generated a list of such genes along with their KEGG ^79^ ortholog (KO) numbers based on existing literature evidence (Supplementary Table S5). We then annotated all isolate genomes using PROKKA v1.14.6 ^64^. The genes resulting from PROKKA gene calling were annotated with EggNOG-mapper v2.1.3 ^80^ to assign KO terms using Diamond ^81^ with 0.0001 e-value and 20.0% identity cut-off values. The resulting KO terms were then searched against our manually curated list of genes. For visualization, the number of KO terms identified in each isolate was normalized by the number of KO terms in the given functional category and a heatmap was created in R using ggplot2 ^49^.

#### DSMZ isolate bundle selection

While all individual collection isolates are available for purchase through DSMZ, we additionally created three cost-efficient bundles of isolates to improve accessibility to the research community (Supplementary Table S6). Bundle A includes 15 representative isolates from the “core” genera that represented 97% of the relative abundance in Swift *et al.* ^41^ (Supplementary Table S3; Fig. 2). Bundle B includes the seven isolates in the SynCom developed by Niu *et al.* ^9^. Bundle C includes 22 representative BSL-1 isolates from all remaining genera including in ZeaMiC that are not represented in Bundles A and B.

**Figure 2.**
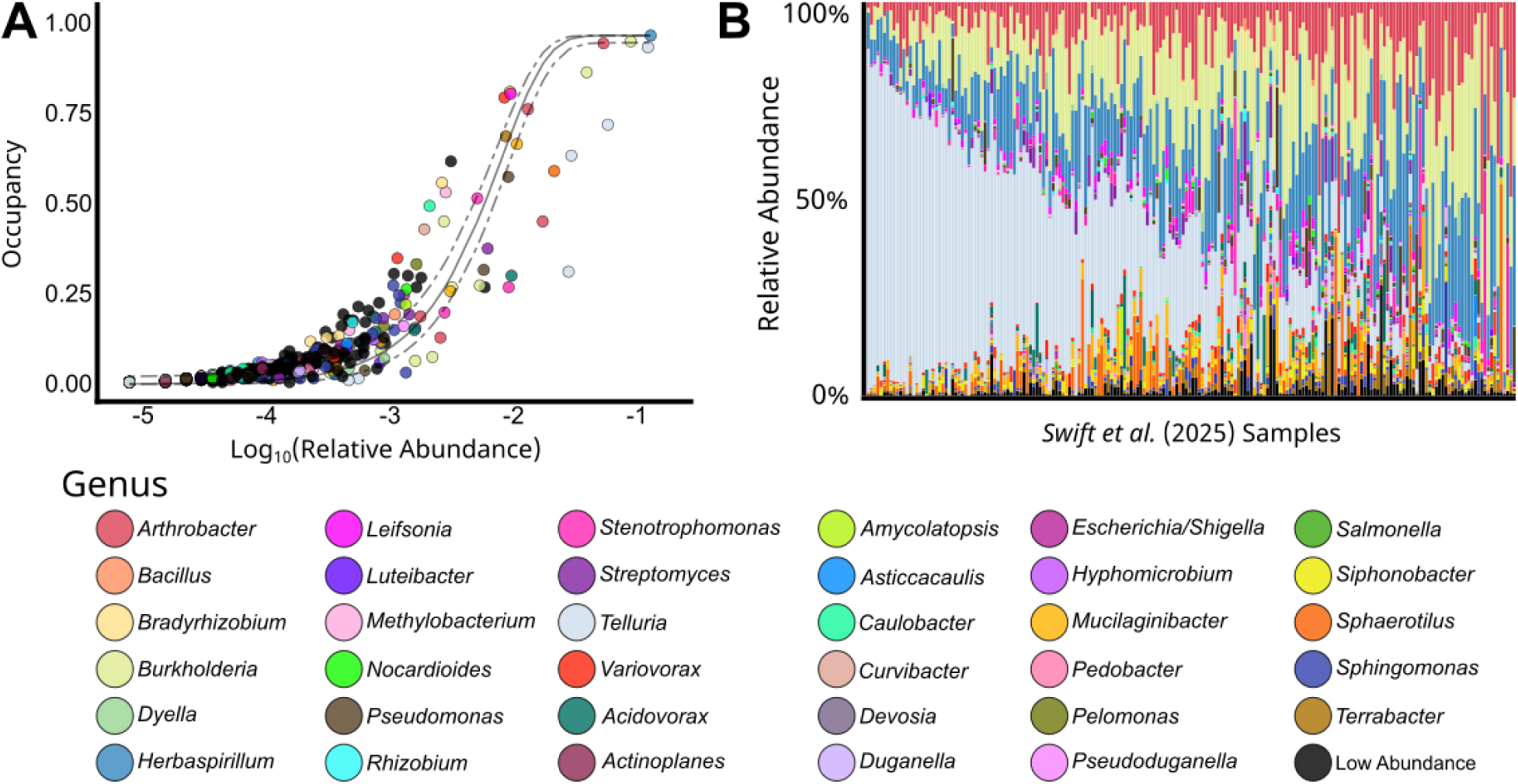
Core genera of a representative maize root microbiome. Briefly, *16S rRNA* gene amplicon sequencing was used to characterize the bacterial root microbiomes of 200 maize plants inoculated with Kansas prairie soil microbiomes (see *Defining core maize root bacterial taxa)*. **A)** Abundance-occupancy curve and **B)** taxonomic barplots for the maize root microbiomes reported in Swift *et al.* 2025 ^41^. A fit neutral model ^82^ to the abundance-occupancy distribution to assess the processes contributing to the microbiome assembly ^42^. Within the taxonomic barplots, each stacked bar represents an individual sample and all genera below the top 35 are condensed into the “Low Abundance” category for visual clarity (n = 98 genera collapsed).

Isolates in Bundles A and C were selected based on their biosafety level categorization (all are biosafety level 1; BSL-1) and genome quality. For genera that are represented by multiple isolates, we selected one representative isolate to include in the bundle based on the number of contigs, with preference being given to isolates with the most contiguous genomes.

The overview of the collection at the DSMZ can be accessed here: https://www.dsmz.de/collection/catalogue/microorganisms/microbiota/zeamic.

## Results

### Dual cultivation strategies recovered ecologically defined core maize root bacteria

To enrich for maize-adapted root colonizers, maize seedlings were grown as bait plants in soil microbiota derived from prairie, post-agricultural, and agricultural soils. Using complementary high-throughput microarray–based and spread plate–based cultivation approaches, we recovered over 900 bacterial isolates associated with maize roots. Genus-level identification of 971 isolates using partial *16S rRNA* gene Sanger sequencing resulted in an initial “crude” maize root bacterial collection (Supplementary Fig. S1-S2 and Table S2). We then curated this “crude” isolate collection to achieve our two design objectives: (i) maximizing overall phylogenetic and taxonomic diversity, and (ii) maximizing representation of core bacterial taxa.

Phylogenetic analysis of partial *16S rRNA* gene sequences from the 971 isolates revealed representation across seven bacterial classes (Supplementary Fig. S1-S2). The collection was strongly dominated by members of the phylum *Pseudomonadota*, with *Gammaproteobacteria*, *Betaproteobacteria*, and *Alphaproteobacteria* together comprising 97.2% of all isolates (946 strains). We used the *16S rRNA* gene tree to guide our selection of candidate strains, pruning redundant isolates while maintaining taxonomic and phylogenetic breadth.

In parallel, we identified “core” taxa to prioritize for inclusion in the curated collection, using existing *16S-V4 rRNA* gene sequencing data from the roots of 200 maize plants inoculated with microbiota from US Midwest soils ^41^. Thus, this study captures maize root bacterial communities across a large number of plants, allowing for identification of prevalent and abundant taxa. Furthermore, the root microbiota measured in that pre-existing study were derived from the same soils used to generate our crude isolate collection, increasing the data’s usefulness for guiding our selection of representative taxa. Following an abundance-occupancy framework ^42^, we defined core bacterial genera as those that had a high relative abundance and were detected in a high proportion of plants (*i.e.*, had high occupancy) in the root (Fig. 2A). Using this approach we identified 35 “core” bacterial genera, which represented 97% (0.13%-31% per genus) of the relative abundance for the total bacterial community and ranged in occupancy between 19-100% of samples (Supplementary Table S3; Fig. 2B). These genera were designated as high-priority targets for inclusion in ZeaMiC. Eight of the 35 core genera were already represented in our crude culture collection, including *Burkholderia, Pseudomonas, Stenotrophomonas, Variovorax, Leifsonia, Bradyrhizobium, Rhizobium*, and *Dyella*. Collectively, these genera accounted for a cumulative mean relative abundance of 25% and exhibited a median occupancy of 83.1% across plants (Fig. 2; Supplementary Table S3), indicating successful recovery of several dominant and persistent maize root microbiome members.

Using these phylogenetic and ecological selection criteria, we advanced 101 “candidate” strains from the crude culture collection for closer evaluation, including whole-genome sequencing. After whole-genome sequencing, we de-replicated closely related isolates that clustered above 98% average nucleotide identity based on whole-genome sequence data, resulting in a curated set of 66 strains representing 26 genera. These included both core and non-core taxa to meet our goals of targeting core bacterial members and maximizing phylogenetic diversity.

Finally, to close the gap of limited coverage of high-priority taxa in the de-replicated US Midwest collection and to capture the additional maize-associated diversity reported across previous studies ^8–17^, we sourced isolates of missing core genera from the global maize–microbiome research community. This included 20 previously isolated maize-associated strains spanning multiple genera, including *Agrobacterium, Arthrobacter, Bacillus, Brucella, Chryseobacterium, Curtobacterium, Deinococcus, Franconibacter, Herbaspirillum, Methylobacterium, Nocardioides, Paenarthrobacter, Pantoea, Peribacillus, Sinomonas, Sphingobium,* and *Streptomyces*. In parallel, we successfully isolated two strains of the genus *Telluria* from maize plants grown with Konza Prairie soil microbiota. *Telluria* was a particularly important target genus due to its dominance (31.6% mean relative abundance, 100% occupancy; Fig. 2B) ^41^ and its close phylogenetic relationship to *Massilia*, a genus with documented effects on maize phenotypes ^59,60^. Together, these efforts added 22 strains to the collection, resulting in a final curated set of 88 maize-associated bacterial isolates (Fig. 3).

**Figure 3.**
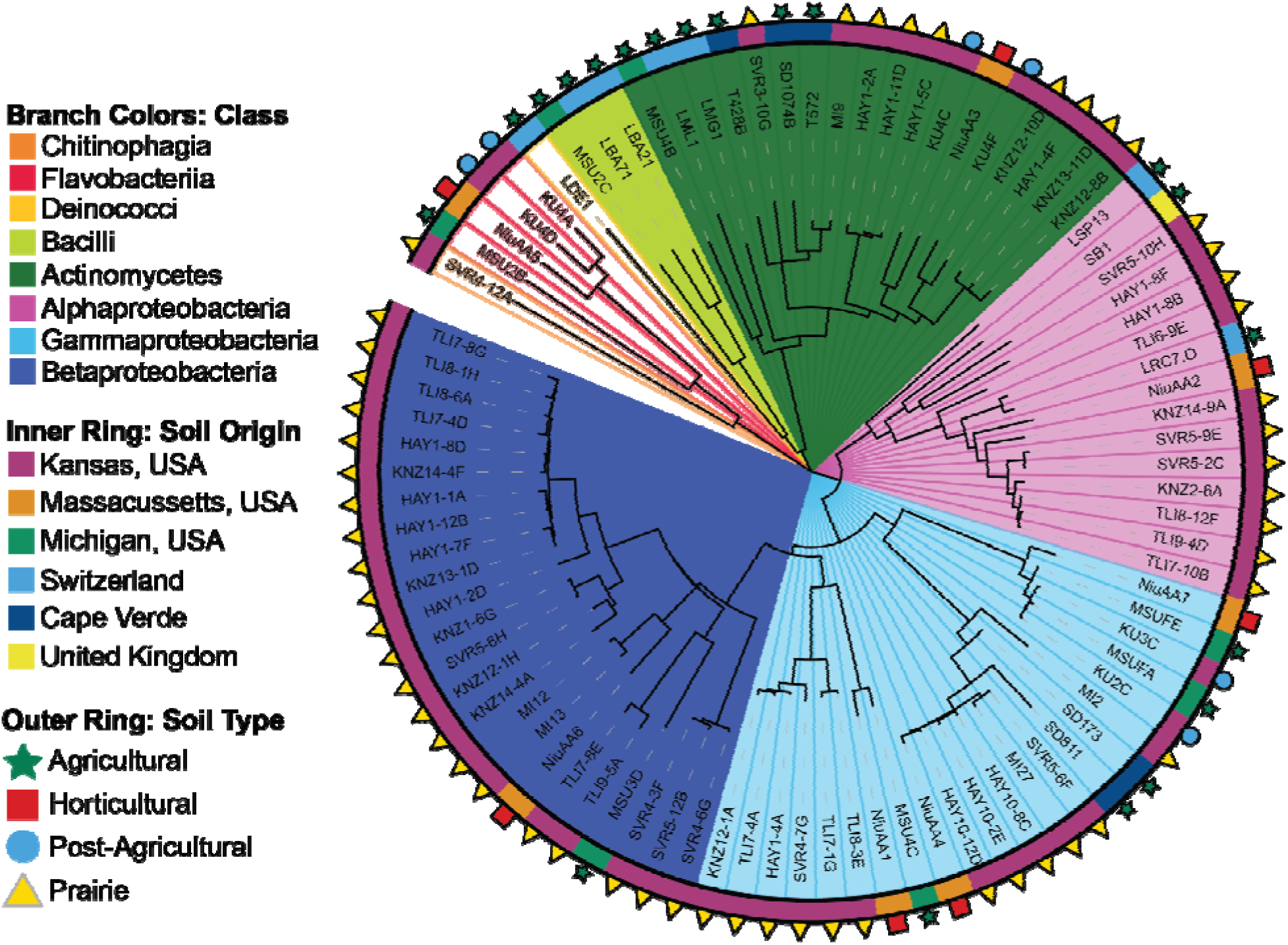
Maximum-likelihood full-length *16S rRNA* gene tree of all 88 isolates in the culture collection. Branches have been colored to indicate taxonomic class. The inner and outer rings indicate the origin and type of soils from which each of the isolates were collected.

### Global prevalence of ZeaMiC isolates across maize root microbiome studies

To assess the global prevalence of ZeaMiC, we analyzed published maize root microbiome studies spanning diverse geographical regions and genotypes, and quantified the prevalence and relative abundance of collection taxa across studies. In addition to Swift *et al.* 2025 ^41^, we aligned our isolates to 16S *rRNA* ASVs from nine other maize microbiome studies conducted in North America and Europe ^12,16–18,60,69–72^ and found a high degree of taxonomic overlap (Fig. 4; Supplementary Table 7). Across the ten studies, between 38 and 83 ZeaMiC isolates were detected in each study at a ≥99% sequence similarity threshold, with detection increasing to between 59 and 87 isolates when the threshold was relaxed to ≥97% sequence similarity. The only ZeaMiC isolate not present in any study was MSUFE, *Pseudomonas koreensis*, however closely related isolates of the same species were recovered across studies (KU3C and MSUFA; Fig. S3&5). This indicates that although ZeaMiC was initially derived from US Midwest soil sources and targeted additions from global collections, we find that its members are abundant across studies and cosmopolitan, both geographically and across different maize compartments.

**Figure 4.**
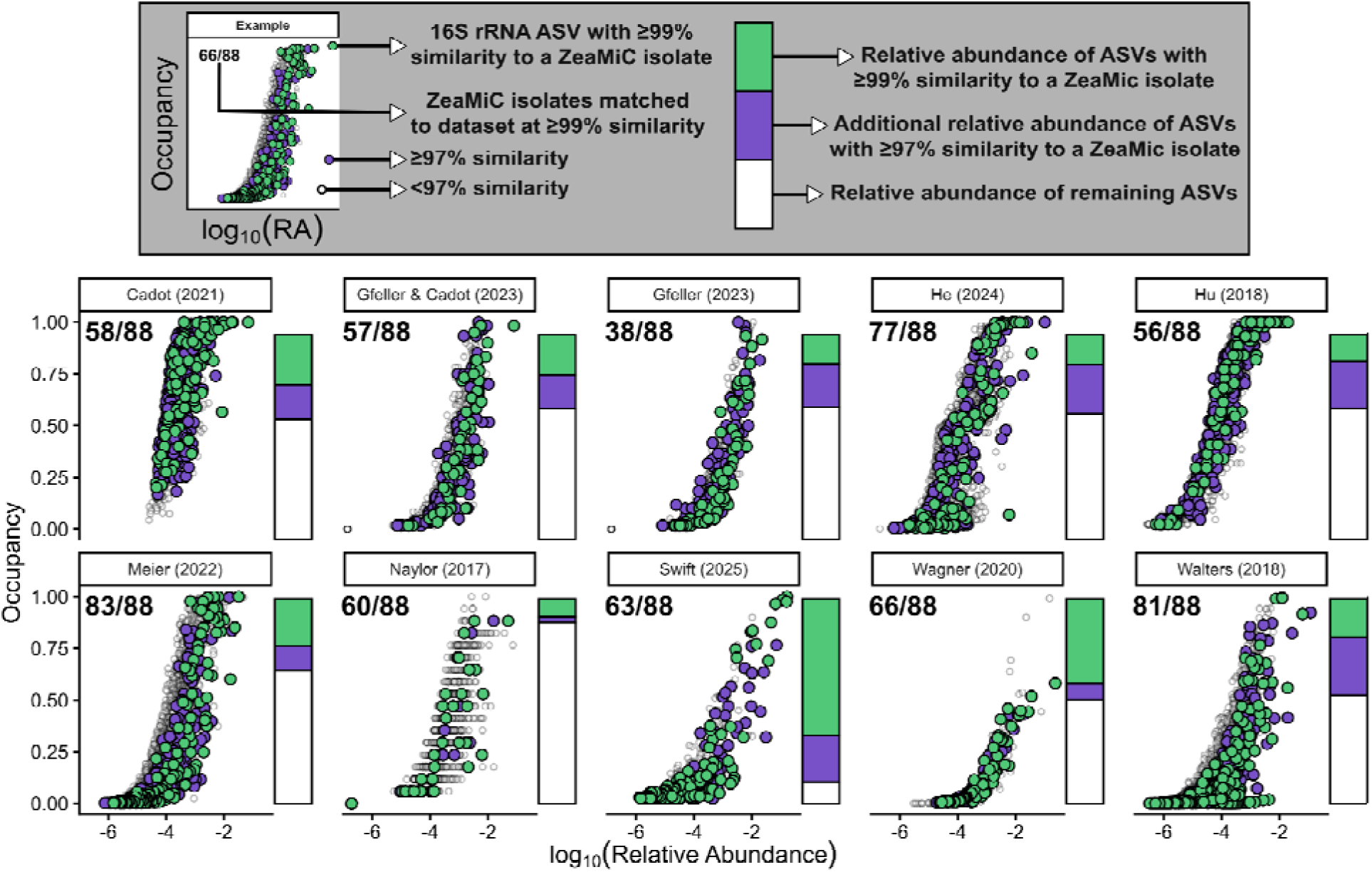
Robust detection of ZeaMiC taxa across diverse maize microbiome studies. For each study (represented by an individual panel), an occupancy vs. abundance curve depicts ASVs with significant alignments to *ZeaMiC* isolates at ≥99% and ≥97% sequence similarit thresholds (green and purple points, respectively). At the top left of each plot, the number of *ZeaMiC* isolates (out of the 88 in the collection) recovered at ≥99% sequence similarity is provided. The stacked barplot to the left depicts the mean relative abundance captured from ASVs with ≥99% sequence similarity to an isolate in *ZeaMiC*. ASVs with ≥97% sequence similarity represent the additional relative abundance captured when relaxing the threshold. Presence and abundance values presented here are drawn from all maize samples across compartments and experimental treatments in each study which provides a more generalized and robust assessment of *ZeaMiC*. Presence and abundance values stratified by compartment are provided in Supplementary Figures S3-6 and Table S8.

Next, we quantified the cumulative relative abundance of ASVs aligning with ZeaMiC taxa within each study. ASVs with ≥99% sequence similarity to ZeaMiC isolates accounted for an average of 23.6% of the total bacterial community relative abundance, ranging from 9.3 to 65.6% across studies. When the sequence similarity threshold was relaxed to ≥97%, this coverage increased to an average of 41% relative abundance (12.1 to 88.9%). Since eight of the ten studies profiled multiple plant compartments, we also evaluated ZeaMiC coverage across compartments (e.g., phyllosphere, rhizosphere, bulk soil) and observed similar patterns (Supplementary Table S8). Considering that soils harbor the most diverse microbial communities in the world, including an estimated 50% of all bacterial species ^83^, the detection of 88 soil-derived isolates across diverse maize datasets at mean relative abundances of 23.6% (≥99% similarity) and 41% (≥97% similarity) is substantial. These findings underscore the consistent recruitment of these taxa by maize and highlight their potential importance within the maize microbiome.

Additionally, we pooled the ASVs from all ten studies, yielding a total of 30,948 ASVs of which 1,193 had ≥99% sequence similarity with ZeaMiC isolates (Fig. 5A). ASVs were pooled and collapsed by genera rather than clustered at a fixed threshold (i.e., 99% or 97% OTUs) as studies utilized different marker regions (e.g., V3–V4, V4, and V5–V7) and resolution. ASVs that aligned to ZeaMiC taxa were broadly distributed across the abundance-occupancy curve, with many of the most prevalent and abundant ASVs represented in our collection (Fig. 5A). Among the top genera represented in ZeaMiC were *Pantoea*, *Arthrobacter*, *Enterobacter*, *Streptomyces*, and *Sphingobium* all of which were recovered in >9 studies (Fig. 5B). Among genera absent from the collection, several notable genera emerged as future targets based on their high relative abundance and prevalence across ≥ 9 studies (Fig. 5B). *Ralstonia* showed the highest mean abundance, with 32 ASVs observed across all studies and a mean relative abundance of 1% (Fig. 5B; Supplementary Table S9). This was the highest relative abundance across all genera (Supplementary Table S9). *Sphingomonas* had the largest number of ASVs with 718 across the 10 studies and a mean relative abundance of 0.03%. Other notable genera included *Terrabacter*, *Amycolatopsis*, *Kribbella*, *Blastococcus*, *Nitrospira*, *Asticcacaulis*, *Lysobacter*, and *Phycicoccus.* These represent future targets for inclusion in ZeaMiC.

**Figure 5.**
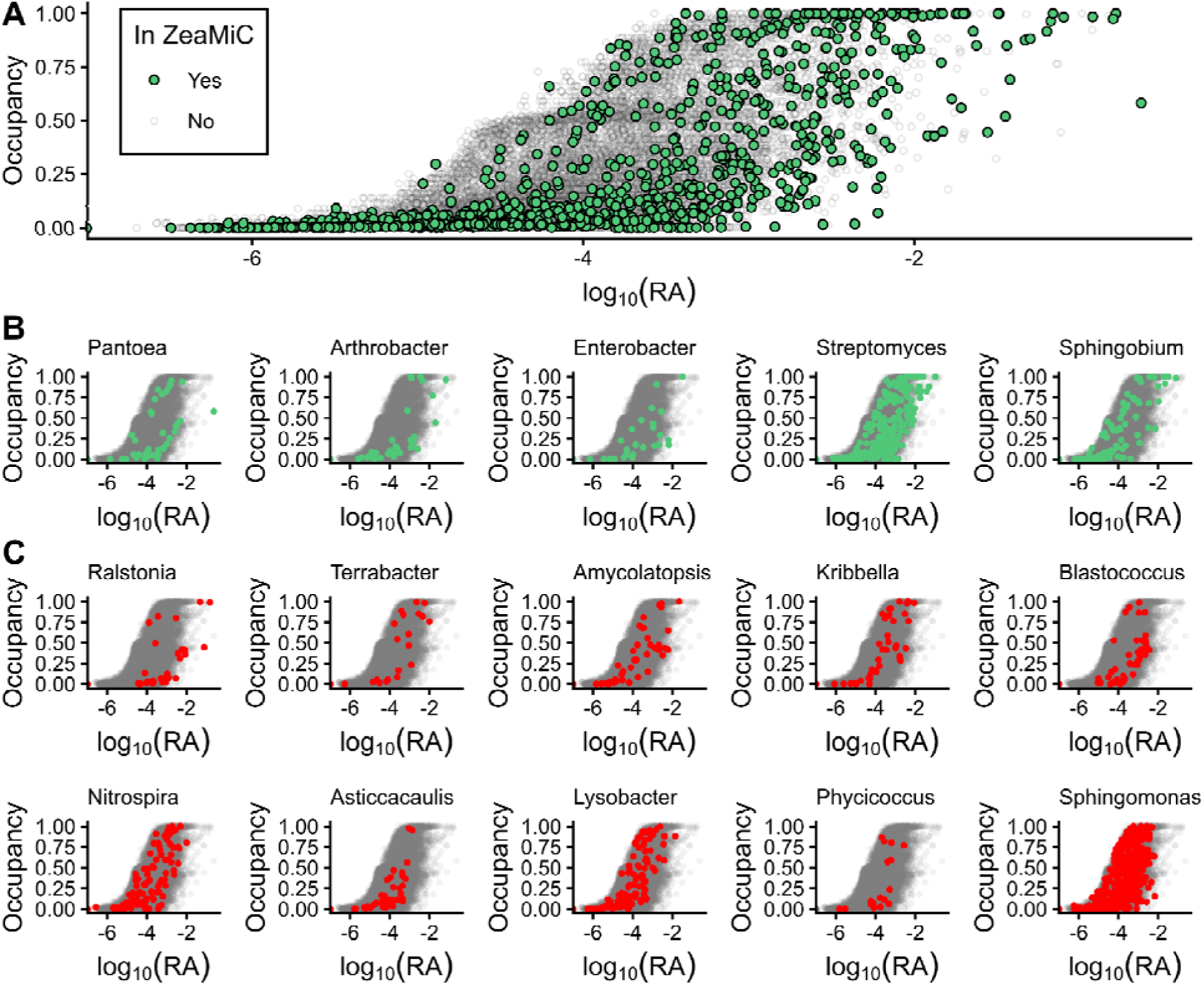
ZeaMiC spans taxa across the full occupancy–abundance space observed among 10 maize microbiome studies. **A)** Occupancy–abundance relationship for ASVs pooled across ten published studies (see Supplementary Table S7). ASVs from each study showing significant alignment at 99% sequence similarity are classified as present in ZeaMiC (green; n = 1,193), whereas all others are classified as absent (grey; n = 29,755 ASVs). **B)** Top genera represented in ZeaMiC and **C)** genera contributing the greatest relative abundance across studies that are not currently represented in ZeaMiC, defined as being present in at least 9 of 10 studies. Within each plot, green or red points depict ASVs recovered from the genus shown in the plot title, while gray points represent all other ASVs. Additional statistics and genera both present and absent from ZeaMiC meeting the studies 9/10 threshold are presented in Supplementary Table S9.

### ZeaMiC genomic diversity and functional potential

To assess the functional potential of our final 88-member isolate collection, we investigated the presence of genes known to contribute to rhizosphere and root colonization ^73–77^, plant growth promotion ^74,75,78^, and nutrient cycling ^76,77^ (Fig. 6, Table S5). Specifically, we annotated isolate genomes for KEGG Orthology (KO) terms associated with biofilm formation, chemotaxis, plant hormone modulation, nitrogen fixation, phosphorus solubilization, and other rhizosphere-relevant processes.

**Figure 6.**
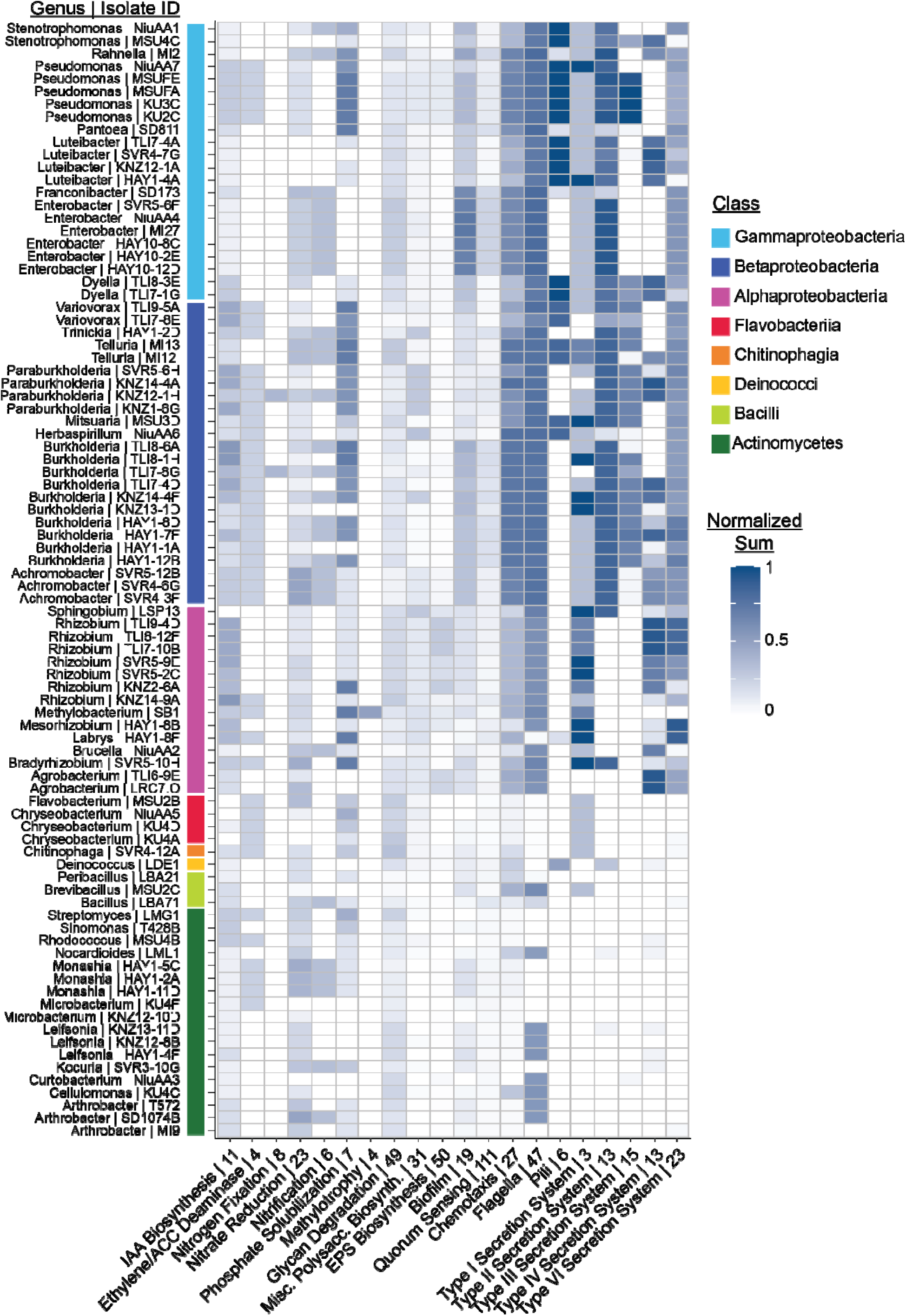
Presence of rhizosphere- and root-colonization, plant growth promotion, and nutrient cycling genes in the collection isolates. Heatmap showing the distribution of KO terms associated with root colonization, plant growth promotion, and nutrient cycling across the 88 maize root-associated bacterial isolates (See Supplementary Table S5 for the full breakdown). For each functional category, gene counts were normalized to the total number of KO terms in that category. The number of KO terms in each category are listed after the functional category name on the x-axis.

The number of KO terms varied substantially across functional categories, therefore gene counts were normalized for each isolate by dividing by the total number of KO terms in that category to generate a proportion. For some functions, such as methylotrophy or nitrogen fixation, the presence of a single key gene might be sufficient to confer the function (e.g. methanol dehydrogenase or nitrogenase, respectively) ^84,85^. In contrast, for multi-component systems such as flagella or the Type VI Secretion System, nearly all associated genes are typically required for full functionality ^86,87^. Thus, putative functions based on genomic evidence should be experimentally validated.

Genes related to chemotaxis, motility, and secretion systems were highly represented across the collection, particularly among Gram-negative isolates (Fig. 6). Similarly, biofilm biosynthesis genes were identified in most isolates (93%). This is consistent with previous findings that these functions play important roles in root colonization and persistence ^73,88–92^. We additionally found degradation genes for plant-derived glycans, such as cellulose, hemicellulose, and pectin, in every isolate. This suggests that plant-derived polysaccharides are an important substrate for many of the isolates and that there may be a wide range of substrate preference and versatility across the collection.

Among plant hormone modulating functions, we found that the ethylene biosynthesis and 1-aminocyclopropane-1-carboxylate (ACC) deaminase pathway was particularly conserved across the Betaproteobacteria, Flavobacteriia, and *Pseudomonas* isolates, though it was also present in several Actinomycetes and Alphaproteobacteria isolates. At least one indole-3-acetic acid (IAA) biosynthesis enzyme was present in 84 of the 88 isolates; however, as with all predicted functions, experimental validation is required to determine whether these isolates are truly capable of synthesizing IAA.

Phosphate solubilization genes were present in 67 of the 88 isolates but were particularly enriched in the *Pseudomonas* isolates and in a number of Alpha- and Betaproteobacteria isolates. We found nitrogen fixation genes (i.e. nitrogenases) encoded in only two isolates – TLI7-8G (*Burkholderia*) and KNZ12-1H (*Paraburkholderia*) – and absent from all *Bradyrhizobium, Rhizobium,* or *Agrobacterium* isolates, even though many members of these three genera are known nitrogen fixers.

## Discussion

In this study, we present a new, publicly available collection of maize root-associated bacteria with whole genomes that is taxonomically representative of the US top corn production region microbiomes. This collection captures most of the phylogenetic diversity of bacterial taxa typically associated with maize roots, and represents isolates collected from a variety of soils (tallgrass and shortgrass prairies, agricultural fields, and post-agricultural fields), geographical locations (US, UK, Switzerland, Cape Verde), and maize genotypes (B73, “Sugar Buns”, and Zm523). This collection includes and expands upon a previously published seven-member Niu SynCom ^9^ and also includes isolates from the collections described by Thoenen *et al.* ^33^ and Custódio *et al.* ^38^. Notably, we have made this resource publicly available for use through DSMZ (live isolates), where all isolates are available for purchase individually and select isolates are available in three cost-efficient bundles (Supplementary Table S6). Whole genomes are additionally available through NCBI.

We recognize that this collection has limitations and reflects the inherent biases of culture-dependent approaches. A few commonly detected and abundant taxa - *Ralstonia*, *Sphingomonas, Terrabacter*, *Amycolatopsis*, *Kribbella*, *Blastococcus*, *Nitrospira*, *Asticcacaulis*, *Lysobacter*, and *Phycicoccus* - are currently absent from our collection, but may play important roles in the maize root microbiome. For instance, *Ralstonia* is commonly studied in agricultural systems due to its pathogenicity to many solanaceous crops ^93,94^, however, we detected this genus in 9 of 10 maize-associated microbiome studies. In two of the studies ^12,16^, *Ralstonia* was the most abundant genus, contributing 5-14% of bacterial relative abundance, and was recovered from every sample (n = 903 and n =3285, respectively). This pattern may reflect the promiscuous nature of the genus, which also has been noted as a contaminant in laboratory reagents and water systems ^95–97^. However, multiple ASVs were recovered for *Ralstonia* within each of these studies and the abundance within these studies was considerable, indicating that these previous detections are unlikely to be driven purely by contamination. Concurrently, a recent genome resource recovered and sequenced a *Ralstonia* sp. (*Ralstonia* sp. R-29, SAMN47200783) associated with the maize rhizosphere ^98^. ZeaMiC includes soil-derived microbes from three land use types, but future work could enhance this collection by systematically isolating from more soil types, different plant compartments (e.g. leaves), and various stages of the plant life cycle (e.g., emergent, juvenile, flowering, grain-filling, etc.) to increase strain diversity. Finally, we note that ZeaMiC is currently solely focused on bacteria. Non-bacterial root-associated microbes, especially archaea, fungi, and protists, will be needed as maize SynCom-based research advances toward higher levels of complexity.

Looking forward, this culture collection will support a wide range of future work investigating maize root-associated microbiota, including mechanistic studies of microbiota-mediated drought tolerance, nutrient acquisition, and plant colonization. More broadly, microbial culture collections are proving to be powerful tools not only for applied studies, but also for testing fundamental ecological principles, such as community assembly, effects of order of arrival on community structure ^99^, and the roles of competition and cooperation in community interactions ^100^, as well as serving as valuable resources for bioactive secondary metabolite discovery ^30^.

The agricultural industry is facing unprecedented challenges due to changing climates, pest and pathogen chemical resistance, and depletion of soil nutrients, among others ^101–104^. Plant-associated microbiomes have the potential to enhance plant growth and resilience ^75,105^, yet challenges remain in developing successful microbiome-informed applications and management strategies ^105–107^. While other, maize microbiome collections have been introduced in recent publications ^9,32,33,38,98^, ZeaMiC is the first globally validated, publicly accessible maize root bacterial isolate collection designed to represent the maize microbiome in the Corn Belt of the US. This diverse 88-member ZeaMiC bacterial culture collection aims to ignite a new phase of functional maize-microbiome studies, provide a reference for ongoing work, and enhance reproducibility. Indeed, this collection will support studies that aim to define causal maize-microbiome relationships and pave new avenues to develop real-world applications for growers.

## Acknowledgments

We thank the staff of the NC State University Phytotron for the use of their facilities and support, Martel Ellis for assisting with initial identification of bacterial isolates, Valéria Custódio for managing and handling the Cape Verde and United Kingdom isolates, Kayla Clouse for providing the MSU soils for use in this project, Martin Chilvers for granting access to the MSU Plant Pathology Research Farm, Mitch Greer, Michaela VonLintel, Justin Roemer, Alan Schlegel, and The Nature Conservancy for granting access to prairie soil collection sites in Kansas, Mirjam Pasch for support in wet lab work for genome sequencing, and the curators and technicians of the German Collection of Microorganisms and Cell Cultures (DSMZ) for valuable discussions during preservation and authentication.

## Author Contribution

MRW and MK designed and supervised the project. AKG, NG, JFS, FT, NEF, and GP cultivated and isolated bacterial strains for the collection and performed *16S rRNA* gene sequencing. AZ performed genome sequencing, assembly, and annotation. AKG, NG, JFS, and CT performed bioinformatic analyses. BN, GC, KS, and CP provided additional bacterial isolates to supplement the collection. RLH provided DSMZ submission guidance and support. AKG, NG, JFS wrote the first draft of the manuscript. All authors provided feedback on the manuscript.

## References

1. Panke-Buisse, K., Poole, A. C., Goodrich, J. K., Ley, R. E. & Kao-Kniffin, J. Selection on soil microbiomes reveals reproducible impacts on plant function. ISME J. 9, 980–989 (2015).

2. Poudel, M. et al. The Role of Plant-Associated Bacteria, Fungi, and Viruses in Drought Stress Mitigation. Front. Microbiol. 12, (2021).

3. Adesemoye, A. O. & Kloepper, J. W. Plant–microbes interactions in enhanced fertilizer-use efficiency. Appl. Microbiol. Biotechnol. 85, 1–12 (2009).

4. Yang, J., Kloepper, J. W. & Ryu, C.-M. Rhizosphere bacteria help plants tolerate abiotic stress. Trends Plant Sci. 14, 1–4 (2009).

5. World Food and Agriculture – Statistical Yearbook. (FAO, 2023). doi:10.4060/cc8166en.

6. Erenstein, O., Jaleta, M., Sonder, K., Mottaleb, K. & Prasanna, B. M. Global maize production, consumption and trade: trends and R&D implications. Food Secur. 14, 1295–1319 (2022).

7. Peiffer, J. A. et al. Diversity and heritability of the maize rhizosphere microbiome under field conditions. Proc. Natl. Acad. Sci. 110, 6548–6553 (2013).

8. Navarro-Noya, Y. E. et al. Bacterial Communities in the Rhizosphere at Different Growth Stages of Maize Cultivated in Soil Under Conventional and Conservation Agricultural Practices. Microbiol. Spectr. 10, e01834–21 (2022).

9. Niu, B., Paulson, J. N., Zheng, X. & Kolter, R. Simplified and representative bacterial community of maize roots. Proc. Natl. Acad. Sci. 114, E2450–E2459 (2017).

10. Guevara-Hernandez, E. et al. Drought induces substitution of bacteria within taxonomic groups in the rhizosphere of native maize from arid and tropical regions. Rhizosphere 29, 100835 (2024).

11. Lopes, L. D. & Schachtman, D. P. Rhizosphere and bulk soil bacterial community succession is influenced more by changes in soil properties than in rhizosphere metabolites across a maize growing season. Appl. Soil Ecol. 189, 104960 (2023).

12. Meier, M. A. et al. Association analyses of host genetics, root-colonizing microbes, and plant phenotypes under different nitrogen conditions in maize. eLife 11, e75790 (2022).

13. Gomes, E. A. et al. Root-Associated Microbiome of Maize Genotypes with Contrasting Phosphorus Use Efficiency. Phytobiomes J. 2, 129–137 (2018).

14. Lund, M. et al. Rhizosphere bacterial communities differ among traditional maize landraces. *Environ*. DNA 4, 1241–1249 (2022).

15. Yadav, P. et al. Zea mays genotype influences microbial and viral rhizobiome community structure. ISME Commun. 3, 129 (2023).

16. Wagner, M. R., Roberts, J. H., Balint-Kurti, P. & Holland, J. B. Heterosis of leaf and rhizosphere microbiomes in field-grown maize. New Phytol. 228, 1055–1069 (2020).

17. Walters, W. A. et al. Large-scale replicated field study of maize rhizosphere identifies heritable microbes. Proc. Natl. Acad. Sci. 115, 7368–7373 (2018).

18. Cadot, S. et al. Specific and conserved patterns of microbiota-structuring by maize benzoxazinoids in the field. Microbiome 9, 103 (2021).

19. Carini, P. A “Cultural” Renaissance: Genomics Breathes New Life into an Old Craft. mSystems 4, 10.1128/msystems.00092-19 (2019).

20. Northen, T. R. et al. Community standards and future opportunities for synthetic communities in plant–microbiota research. Nat. Microbiol. 9, 2774–2784 (2024).

21. Vorholt, J. A., Vogel, C., Carlström, C. I. & Müller, D. B. Establishing Causality: Opportunities of Synthetic Communities for Plant Microbiome Research. Cell Host Microbe 22, 142–155 (2017).

22. Bai, Y. et al. Functional overlap of the Arabidopsis leaf and root microbiota. Nature 528, 364–369 (2015).

23. Schäfer, M., Vogel, C. M., Bortfeld-Miller, M., Mittelviefhaus, M. & Vorholt, J. A. Mapping phyllosphere microbiota interactions in planta to establish genotype–phenotype relationships. Nat. Microbiol. 7, 856–867 (2022).

24. Vogel, C. M., Potthoff, D. B., Schäfer, M., Barandun, N. & Vorholt, J. A. Protective role of the Arabidopsis leaf microbiota against a bacterial pathogen. Nat. Microbiol. 6, 1537–1548 (2021).

25. Wippel, K. et al. Host preference and invasiveness of commensal bacteria in the Lotus and Arabidopsis root microbiota. Nat. Microbiol. 6, 1150–1162 (2021).

26. Robertson-Albertyn, S. et al. Genome-Annotated Bacterial Collection of the Barley Rhizosphere Microbiota. Microbiol. Resour. Announc. 11, e01064–21 (2022).

27. Armanhi, J. S. L. et al. A Community-Based Culture Collection for Targeting Novel Plant Growth-Promoting Bacteria from the Sugarcane Microbiome. Front. Plant Sci. 8, (2018).

28. Grady, K. L. & Shade, A. Bacterial isolate collection from switchgrass rhizosphere. Microbiol. Resour. Announc. 13, e01133–23 (2024).

29. Bertani, I., Abbruscato, P., Piffanelli, P., Subramoni, S. & Venturi, V. Rice bacterial endophytes: isolation of a collection, identification of beneficial strains and microbiome analysis. Environ. Microbiol. Rep. 8, 388–398 (2016).

30. Blacutt, A. et al. An In Vitro Pipeline for Screening and Selection of Citrus-Associated Microbiota with Potential Anti-“Candidatus Liberibacter asiaticus” Properties. Appl. Environ. Microbiol. 86, e02883–19 (2020).

31. Carper, D. L. et al. Cultivating the Bacterial Microbiota of Populus Roots. mSystems 6, e01306–20 (2021).

32. Dai, R. et al. Crop root bacterial and viral genomes reveal unexplored species and microbiome patterns. Cell 188, 2521–2539.e22 (2025).

33. Thoenen, L., et al. Bacterial tolerance to host-exuded specialized metabolites structures the maize root microbiome. Proc. Natl. Acad. Sci. 120, e2310134120 (2023).

34. Thoenen, L. et al. The lactonase BxdA mediates metabolic specialisation of maize root bacteria to benzoxazinoids. Nat. Commun. 15, 6535 (2024).

35. Thoenen, L. et al. Synthetic communities of maize root bacteria interact and redirect benzoxazinoid metabolization. mSphere 10, e00159–25 (2025).

36. Wagner, M. R. et al. Microbe-dependent heterosis in maize. Proc. Natl. Acad. Sci. 118, e2021965118 (2021).

37. Garrell, A.-K. et al. Differential metaproteomics of bacteria grown in vitro and in planta reveals functions used during growth on maize roots. bioRxiv 2025.06.02.657423 (2025) doi:10.1101/2025.06.02.657423.

38. Custódio, V. et al. Individual leaf microbiota tunes a genetic regulatory network to promote leaf growth. Cell Host Microbe 33, 436–450.e15 (2025).

39. U.S. Department of Agriculture, Foreign Agricultural Service. Corn Production. (2025).

40. Wang, S., Di Tommaso, S., Deines, J. M. & Lobell, D. B. Mapping twenty years of corn and soybean across the US Midwest using the Landsat archive. Sci. Data 7, 307 (2020).

41. Swift, J. F. et al. Drought stress homogenizes maize growth responses to diverse natural soil microbiomes. Plant Soil 509, 181–199 (2025).

42. Shade, A. & Stopnisek, N. Abundance-occupancy distributions to prioritize plant core microbiome membership. Curr. Opin. Microbiol. 49, 50–58 (2019).

43. Tovar-Herrera, O. E. et al. Core rumen microbes are functional generalists that sustain host metabolism and gut ecosystem function. *Nat*. Ecol. Evol. 10, 44–58 (2026).

44. Altschul, S. F., Gish, W., Miller, W., Myers, E. W. & Lipman, D. J. Basic local alignment search tool. J. Mol. Biol. 215, 403–410 (1990).

45. Edgar, R. C. MUSCLE: multiple sequence alignment with high accuracy and high throughput. Nucleic Acids Res. 32, 1792–1797 (2004).

46. Kumar, S., Stecher, G., Li, M., Knyaz, C. & Tamura, K. MEGA X: Molecular Evolutionary Genetics Analysis across Computing Platforms. Mol. Biol. Evol. 35, 1547–1549 (2018).

47. Steenwyk, J. L., Iii, T. J. B., Li, Y., Shen, X.-X. & Rokas, A. ClipKIT: A multiple sequence alignment trimming software for accurate phylogenomic inference. PLOS Biol. 18, e3001007 (2020).

48. Letunic, I. & Bork, P. Interactive Tree Of Life (iTOL) v5: an online tool for phylogenetic tree display and annotation. Nucleic Acids Res. 49, W293–W296 (2021).

49. Wickham, H. Ggplot2: Elegant Graphics for Data Analysis. (Springer-Verlag New York, 2016).

50. Hrovat, K., Dutilh, B. E., Medema, M. H. & Melkonian, C. Taxonomic resolution of different 16S rRNA variable regions varies strongly across plant-associated bacteria. ISME Commun. 4, ycae034 (2024).

51. Blake, C., Christensen, M. N. & Kovács, Á. T. Molecular Aspects of Plant Growth Promotion and Protection by Bacillus subtilis. Mol. Plant-Microbe Interactions® 34, 15–25 (2021).

52. Singh, R. & Goodwin, S. B. Exploring the Corn Microbiome: A Detailed Review on Current Knowledge, Techniques, and Future Directions. PhytoFrontiers^TM^ 2, 158–175 (2022).

53. Vanissa, T. T. G. et al. The Response of Maize to Inoculation with Arthrobacter sp. and Bacillus sp. in Phosphorus-Deficient, Salinity-Affected Soil. Microorganisms 8, 1005 (2020).

54. Stephan, R. et al. Re-examination of the taxonomic status of Enterobacter helveticus, Enterobacter pulveris and Enterobacter turicensis as members of the genus Cronobacter and their reclassification in the genera Franconibacter gen. nov. and Siccibacter gen. nov. as Franconibacter helveticus comb. nov., Franconibacter pulveris comb. nov. and Siccibacter turicensis comb. nov., respectively. Int. J. Syst. Evol. Microbiol. 64, 3402–3410 (2014).

55. Krumbach, J. et al. Metabolic analysis of a bacterial synthetic community from maize roots provides new mechanistic insights into microbiome stability. bioRxiv 2021.11.28.470254 (2021) doi:10.1101/2021.11.28.470254.

56. van Schaik, J., Li, Z., Cheadle, J. & Crook, N. Engineering the Maize Root Microbiome: A Rapid MoClo Toolkit and Identification of Potential Bacterial Chassis for Studying Plant–Microbe Interactions. ACS Synth. Biol. 12, 3030–3040 (2023).

57. Garrell, A.-K., Vintila, S., Phillip, H., Beck, A. E. & Kleiner, M. A Minimal Medium for Culturing Maize Root-Associated Microbes Based on a Plant Growth Medium. bioRxiv 2025.07.07.663485 (2025) doi:10.1101/2025.07.07.663485.

58. Bowman, J. P. Genome-wide and constrained ordination-based analyses of EC code data support reclassification of the species of Massilia La Scola et al. 2000 into Telluria Bowman et al. 1993, Mokoshia gen. nov. and Zemynaea gen. nov. Int. J. Syst. Evol. Microbiol. 73, (2023).

59. Wang, D. et al. Lateral root enriched Massilia associated with plant flowering in maize. Microbiome 12, 124 (2024).

60. He, X. et al. Heritable microbiome variation is correlated with source environment in locally adapted maize varieties. Nat. Plants 10, 598–617 (2024).

61. Parnell, J. J. et al. Effective Seed Sterilization Methods Require Optimization Across Maize Genotypes. Phytobiomes J. PBIOMES-12-23-0137-R (2024) doi:10.1094/PBIOMES-12-23-0137-R.

62. Prjibelski, A., Antipov, D., Meleshko, D., Lapidus, A. & Korobeynikov, A. Using SPAdes De Novo Assembler. Curr. Protoc. Bioinforma. 70, e102 (2020).

63. Gurevich, A., Saveliev, V., Vyahhi, N. & Tesler, G. QUAST: quality assessment tool for genome assemblies. Bioinformatics 29, 1072–1075 (2013).

64. Seemann, T. Prokka: rapid prokaryotic genome annotation. Bioinformatics 30, 2068–2069 (2014).

65. Olm, M. R., Brown, C. T., Brooks, B. & Banfield, J. F. dRep: a tool for fast and accurate genomic comparisons that enables improved genome recovery from metagenomes through de-replication. ISME J. 11, 2864–2868 (2017).

66. Parks, D. H., Imelfort, M., Skennerton, C. T., Hugenholtz, P. & Tyson, G. W. CheckM: assessing the quality of microbial genomes recovered from isolates, single cells, and metagenomes. Genome Res. 25, 1043–1055 (2015).

67. Simão, F. A., Waterhouse, R. M., Ioannidis, P., Kriventseva, E. V. & Zdobnov, E. M. BUSCO: assessing genome assembly and annotation completeness with single-copy orthologs. Bioinforma. Oxf. Engl. 31, 3210–3212 (2015).

68. Kolmogorov, M., Yuan, J., Lin, Y. & Pevzner, P. A. Assembly of long, error-prone reads using repeat graphs. Nat. Biotechnol. 37, 540–546 (2019).

69. Naylor, D., DeGraaf, S., Purdom, E. & Coleman-Derr, D. Drought and host selection influence bacterial community dynamics in the grass root microbiome. ISME J. 11, 2691–2704 (2017).

70. Hu, L. et al. Root exudate metabolites drive plant-soil feedbacks on growth and defense by shaping the rhizosphere microbiota. Nat. Commun. 9, 2738 (2018).

71. Gfeller, V. et al. Plant secondary metabolite-dependent plant-soil feedbacks can improve crop yield in the field. eLife 12, e84988 (2023).

72. Gfeller, V. et al. Soil chemical and microbial gradients determine accumulation of root-exuded secondary metabolites and plant–soil feedbacks in the field. J. Sustain. Agric. Environ. 2, 173–188 (2023).

73. Levy, A. et al. Genomic features of bacterial adaptation to plants. Nat. Genet. 50, 138–150 (2018).

74. Zhang, P. et al. The Distribution of Tryptophan-Dependent Indole-3-Acetic Acid Synthesis Pathways in Bacteria Unraveled by Large-Scale Genomic Analysis. Mol. Basel Switz. 24, 1411 (2019).

75. Compant, S. et al. Harnessing the plant microbiome for sustainable crop production. Nat. Rev. Microbiol. 23, 9–23 (2025).

76. Tu, Q., Lin, L., Cheng, L., Deng, Y. & He, Z. NCycDB: a curated integrative database for fast and accurate metagenomic profiling of nitrogen cycling genes. Bioinforma. Oxf. Engl. 35, 1040–1048 (2019).

77. Zeng, J. et al. PCycDB: a comprehensive and accurate database for fast analysis of phosphorus cycling genes. Microbiome 10, 101 (2022).

78. Bruto, M., Prigent-Combaret, C., Muller, D. & Moënne-Loccoz, Y. Analysis of genes contributing to plant-beneficial functions in Plant Growth-Promoting Rhizobacteria and related Proteobacteria. Sci. Rep. 4, 6261 (2014).

79. Kanehisa, M., Sato, Y., Kawashima, M., Furumichi, M. & Tanabe, M. KEGG as a reference resource for gene and protein annotation. Nucleic Acids Res. 44, D457–D462 (2016).

80. Cantalapiedra, C. P., Hernández-Plaza, A., Letunic, I., Bork, P. & Huerta-Cepas, J. eggNOG-mapper v2: Functional Annotation, Orthology Assignments, and Domain Prediction at the Metagenomic Scale. Mol. Biol. Evol. 38, 5825–5829 (2021).

81. Buchfink, B., Reuter, K. & Drost, H.-G. Sensitive protein alignments at tree-of-life scale using DIAMOND. Nat. Methods 18, 366–368 (2021).

82. Burns, A. R. et al. Contribution of neutral processes to the assembly of gut microbial communities in the zebrafish over host development. ISME J. 10, 655–664 (2016).

83. Anthony, M. A., Bender, S. F. & van der Heijden, M. G. A. Enumerating soil biodiversity. Proc. Natl. Acad. Sci. 120, e2304663120 (2023).

84. Le, T.-K., Lee, Y.-J., Han, G. H. & Yeom, S.-J. Methanol Dehydrogenases as a Key Biocatalysts for Synthetic Methylotrophy. Front. Bioeng. Biotechnol. 9, 787791 (2021).

85. Rucker, H. R. & Kaçar, B. Enigmatic evolution of microbial nitrogen fixation: insights from Earth’s past. Trends Microbiol. 32, 554–564 (2024).

86. Minamino, T. & Kinoshita, M. Structure, Assembly, and Function of Flagella Responsible for Bacterial Locomotion. EcoSal Plus 11, eesp-0011-2023 (2023).

87. Cianfanelli, F. R., Monlezun, L. & Coulthurst, S. J. Aim, Load, Fire: The Type VI Secretion System, a Bacterial Nanoweapon. Trends Microbiol. 24, 51–62 (2016).

88. Bernal, P., Llamas, M. A. & Filloux, A. Type VI secretion systems in plant-associated bacteria. Environ. Microbiol. 20, 1–15 (2018).

89. Büttner, D. & He, S. Y. Type III Protein Secretion in Plant Pathogenic Bacteria. Plant Physiol. 150, 1656–1664 (2009).

90. Knights, H. E., Jorrin, B., Haskett, T. L. & Poole, P. S. Deciphering bacterial mechanisms of root colonization. Environ. Microbiol. Rep. 13, 428–444 (2021).

91. Scharf, B. E., Hynes, M. F. & Alexandre, G. M. Chemotaxis signaling systems in model beneficial plant-bacteria associations. Plant Mol. Biol. 90, 549–559 (2016).

92. de Weert, S. et al. Flagella-driven chemotaxis towards exudate components is an important trait for tomato root colonization by Pseudomonas fluorescens. Mol. Plant-Microbe Interact. MPMI 15, 1173–1180 (2002).

93. Vailleau, F. & Genin, S. Ralstonia solanacearum: An Arsenal of Virulence Strategies and Prospects for Resistance. Annu. Rev. Phytopathol. 61, 25–47 (2023).

94. García, R. O., Kerns, J. P. & Thiessen, L. Ralstonia solanacearum Species Complex: A Quick Diagnostic Guide. Plant Health Prog. 20, 7–13 (2019).

95. Grahn, N., Olofsson, M., Ellnebo-Svedlund, K., Monstein, H.-J. & Jonasson, J. Identification of mixed bacterial DNA contamination in broad-range PCR amplification of 16S rDNA V1 and V3 variable regions by pyrosequencing of cloned amplicons. FEMS Microbiol. Lett. 219, 87–91 (2003).

96. Salter, S. J. et al. Reagent and laboratory contamination can critically impact sequence-based microbiome analyses. BMC Biol. 12, 87 (2014).

97. Laurence, M., Hatzis, C. & Brash, D. E. Common Contaminants in Next-Generation Sequencing That Hinder Discovery of Low-Abundance Microbes. PLOS ONE 9, e97876 (2014).

98. Paulsen, A. A., Vang, D. X. & Halverson, L. J. Complete genome sequences of 37 bacteria in a maize rhizosphere synthetic community. Microbiol. Resour. Announc. 14, e00496–25 (2025).

99. Carlström, C. I. et al. Synthetic microbiota reveal priority effects and keystone strains in the Arabidopsis phyllosphere. *Nat*. Ecol. Evol. 3, 1445–1454 (2019).

100. Schäfer, M. et al. Metabolic interaction models recapitulate leaf microbiota ecology. Science 381, eadf5121 (2023).

101. Gowda, P. et al. Chapter 10: Agriculture and Rural Communities. In Impacts, Risks, and Adaptation in the United States: Fourth National Climate Assessment vol. 2 (U.S. Global Change Research Program, Washington, DC, USA, 2018).

102. Rosenzweig, C. et al. Assessing agricultural risks of climate change in the 21st century in a global gridded crop model intercomparison. Proc. Natl. Acad. Sci. 111, 3268–3273 (2014).

103. Kirchmann, H. & Thorvaldsson, G. Challenging targets for future agriculture. Eur. J. Agron. 12, 145–161 (2000).

104. Geffersa, A. G. et al. The socio-economic challenges of managing pathogen evolution in agriculture. Philos. Trans. R. Soc. B Biol. Sci. 378, 20220012 (2023).

105. Parnell, J. J. et al. From the Lab to the Farm: An Industrial Perspective of Plant Beneficial Microorganisms. Front. Plant Sci. 7, (2016).

106. Copeland, C., Schulze-Lefert, P. & Ma, K.-W. Potential and challenges for application of microbiomes in agriculture. Plant Cell 37, koaf185 (2025).

107. Sessitsch, A., Pfaffenbichler, N. & Mitter, B. Microbiome Applications from Lab to Field: Facing Complexity. Trends Plant Sci. 24, 194–198 (2019).

