## Supplementary Appendix for "ZeaMiC: a Publicly Available Culture Collection of Maize Root-Associated Bacteria"

**This PDF file includes:**

Supplementary Text

Supplementary Figures S1 to S6

Tables S1 to S9

### Supplementary Text

##### ZeaMiC Comparison to Maize Microbiomes

To compare the relevance of the ZeaMiC collection with maize microbiomes from a broad geographic distribution we selected 10 studies from across North America and Europe (Naylor *et al.*, 2017; Walters *et al.*, 2018; Hu *et al.*, 2018; Wagner *et al.*, 2020; Cadot *et al.*, 2021; Meier *et al.*, 2022; Gfeller *et al.*, 2023a,b; He *et al.*, 2024; Swift *et al.*, 2025). These studies each employed amplicon sequencing targeting various regions of the 16S rRNA gene (V3–V4, V4, and V5–V7). For each study, metadata and weblinks were retrieved from the National Center for Biotechnology Information Sequence Read Archive using ffq v0.3.1 (Gálvez-Merchán *et al.*, 2023). For four studies, intermediate files (e.g., ASV tables) were available, which we used directly in the downstream analysis. For the others, we downloaded raw sequence reads in FASTQ format and processed them into ASV tables. The exact parameters used for sequence processing varied among datasets depending on the primers used, read quality, read length, and other factors; however, the core steps were the same. First, we used cutadapt (Martin, 2011) to remove 5’ and 3’ primer sequences, simultaneously discarding reads with an expected error rate greater than 0.15. Next, we used dada2 (Callahan *et al.*, 2016) to estimate error rates from a concatenated file containing 50-200 reads per sample in the dataset. For one dataset (He et al. 2024), error rate estimation failed due to uniformity of quality scores, so we instead used the empirical error matrices that are provided in the dada2 package. Based on visualized read quality using fastQC (http://www.bioinformatics.babraham.ac.uk/projects/fastqc), we chose expected error rate thresholds and truncation lengths that would remove low-quality sequences from each dataset while retaining sufficient data for downstream analysis. Quality filtering, denoising, dereplication, and de novo chimera removal (using the consensus method) were implemented using the dada2 pipeline. Finally, we assigned taxonomy to each ASV using the RDP classifier trained on the RDP training set version 19 supplemented with plant mitochondrial sequences (Wang *et al.*, 2007; Wang & Cole, 2024).

After initial processing, ASV tables were further filtered individually using phyloseq v1.54 (McMurdie & Holmes, 2013) to remove host plastid contamination, samples with low sequence depth, and “non-reproducible” ASVs. ASVs were classified as “non-reproducible” if they did not occur in more than five samples at a depth of at least 25 reads. This filtering substantially reduces the total number of ASVs while retaining more than 90% of the original sequencing reads. From each study, ASV sequences were exported to a fasta file, which were then all concatenated and used to create a searchable sequence database. Database construction and alignments were handled in BLAST+ v2.17.0 (Camacho *et al.*, 2009). We aligned the full-length 16S rRNA gene sequences from the ZeaMiC collection to this database using two identity thresholds: 97% and 99%. Presence was assessed for each ZeaMiC isolate for each dataset, datasets were also assessed individually by which maize microbiome compartment was profiled (Supplementary Figure S3-6). Additionally, we quantified the relative abundance in each dataset contributed by ASVs with significant alignments to ZeaMiC isolates across each compartment profiled (Table S8).

### Supplementary Figures

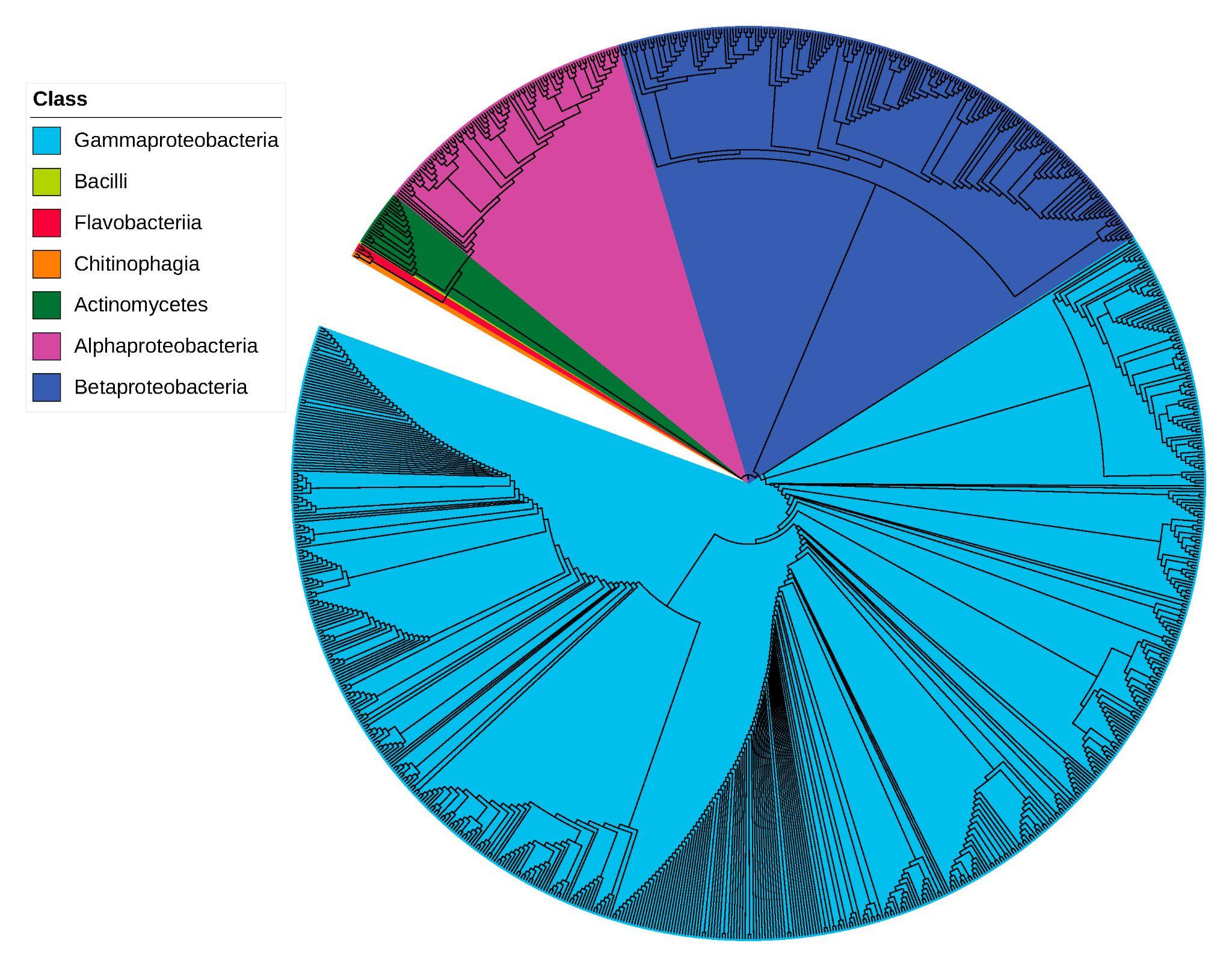

**Figure S1. Crude culture collection partial 16S rRNA gene tree.** The tree includes 971 bacterial isolates that were cultured using array-based and spread plate-based approaches. Details about each isolate including taxonomy, origins, isolation method, and 16S rRNA sequence are provided in Supplementary Table S2. Colors represent bacterial Class.

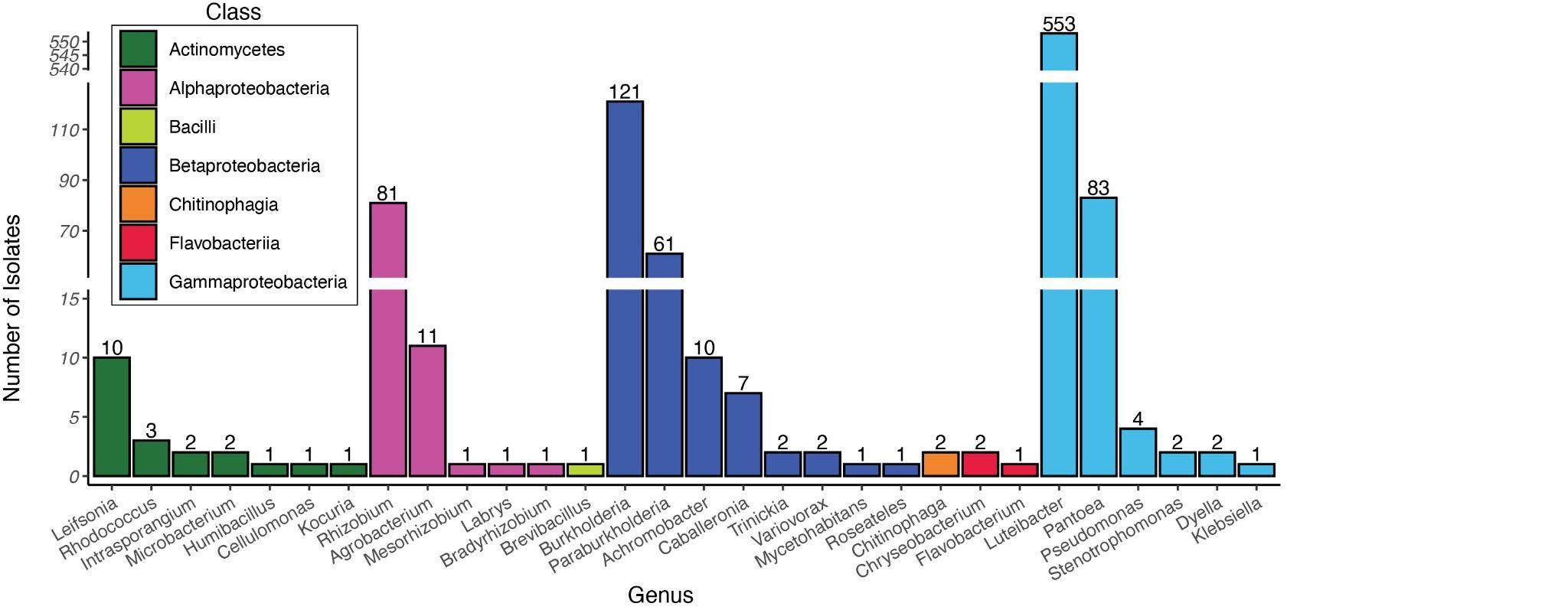

**Figure S2. Crude culture collection isolates counts.** Bar plot indicating the number of US Midwest soil-derived isolates assigned to each putative bacterial genus based on partial *16S rRNA* gene sequencing. Colors correspond to bacterial class.

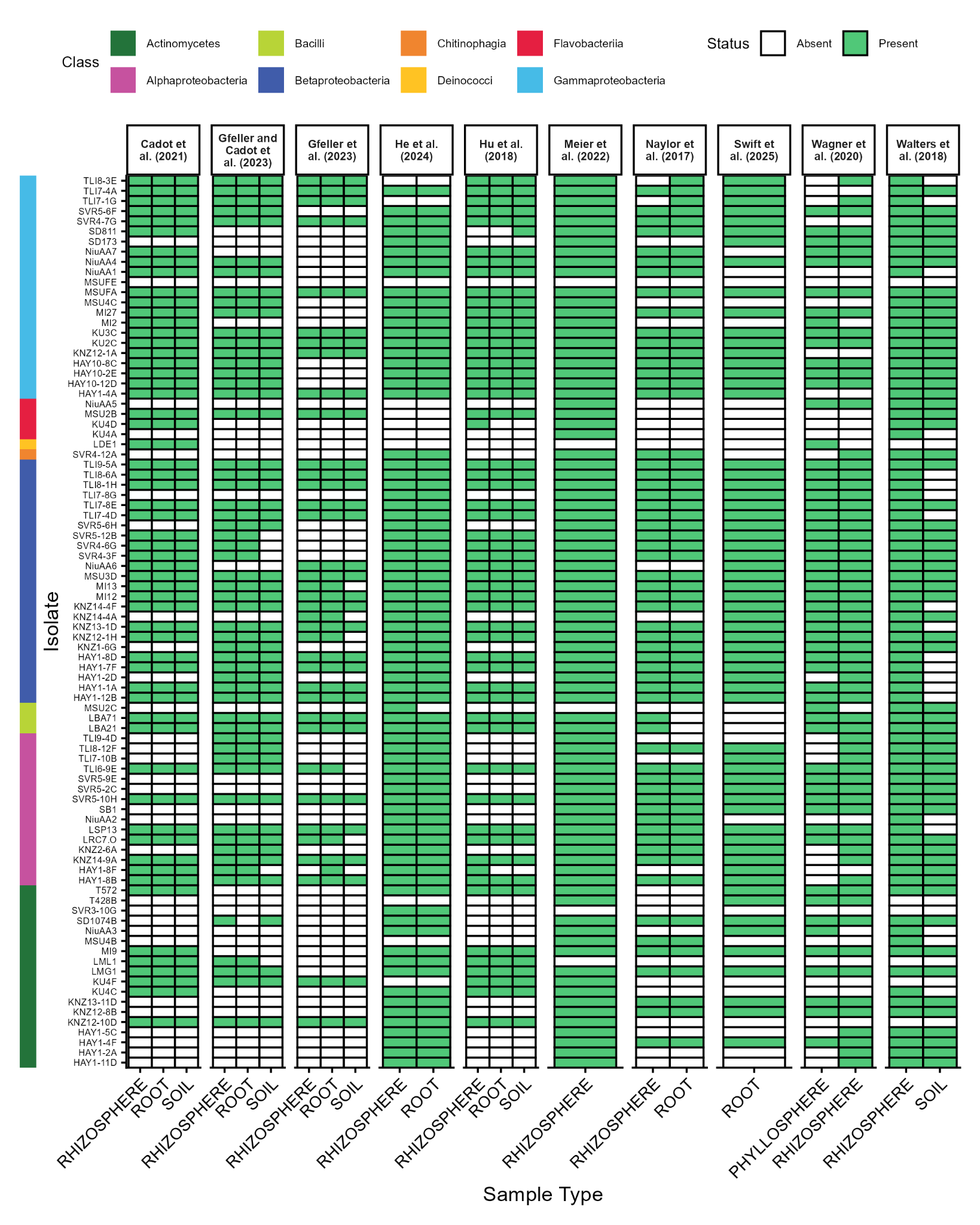

**Figure S3. The presence of ZeaMiC isolates in maize 16S rRNA gene microbiome datasets at 99% identity.** Significant alignments were filtered using a 99% identity threshold. For datasets containing multiple maize microbiome compartments, each compartment is shown as an individual subset within the study panel. ZeaMiC isolate labels are arranged by class (see Table S4 for full taxonomy of each ZeaMiC isolate).

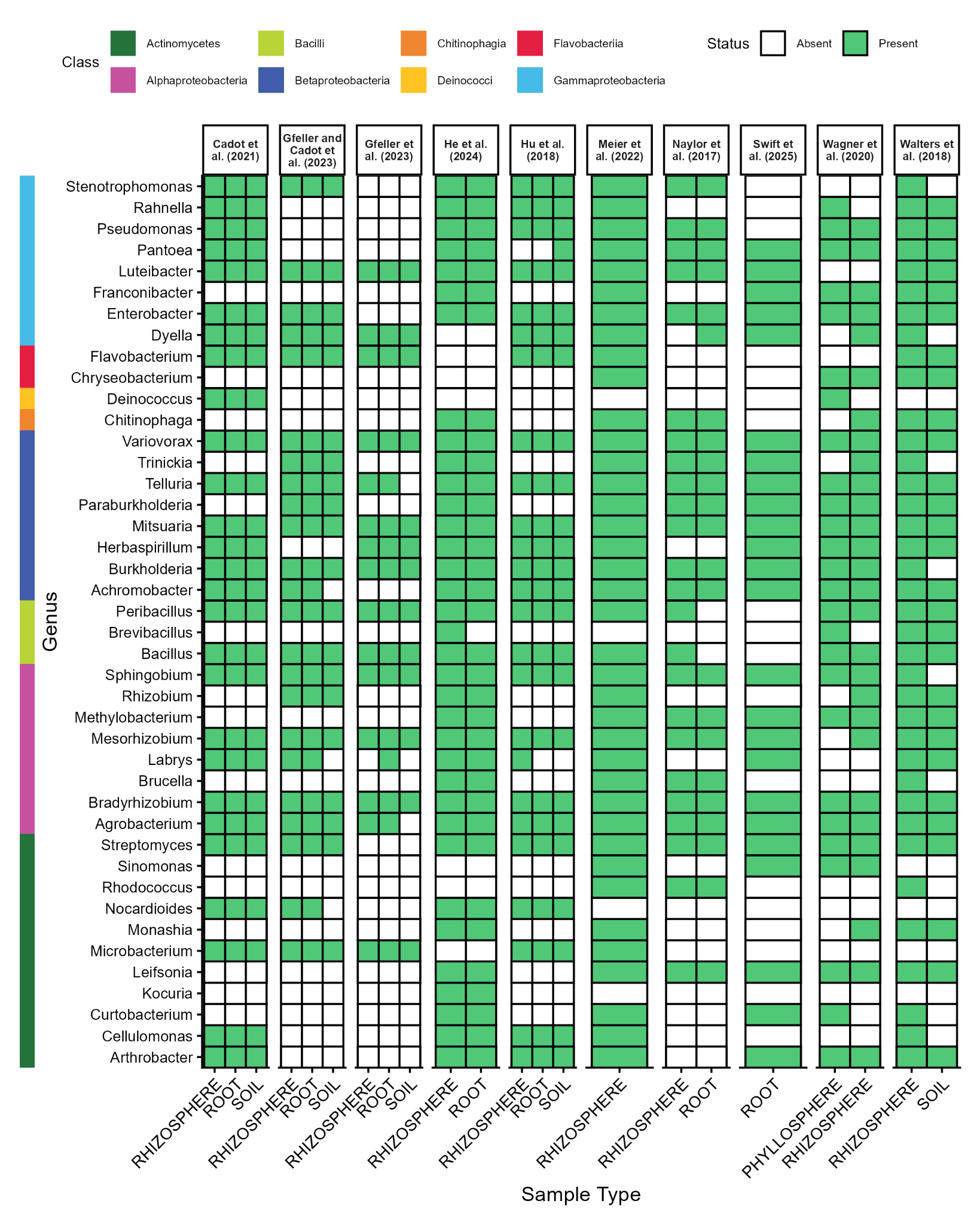

**Figure S4. The presence of ZeaMiC genera in maize 16S rRNA gene microbiome datasets at 99% identity.** Significant alignments were filtered using a 99% identity threshold. For datasets containing multiple maize microbiome compartments, each compartment is shown as an individual subset within the study panel. ZeaMiC genera labels are arranged by class (see Table S4 for full taxonomy of each ZeaMiC isolate).

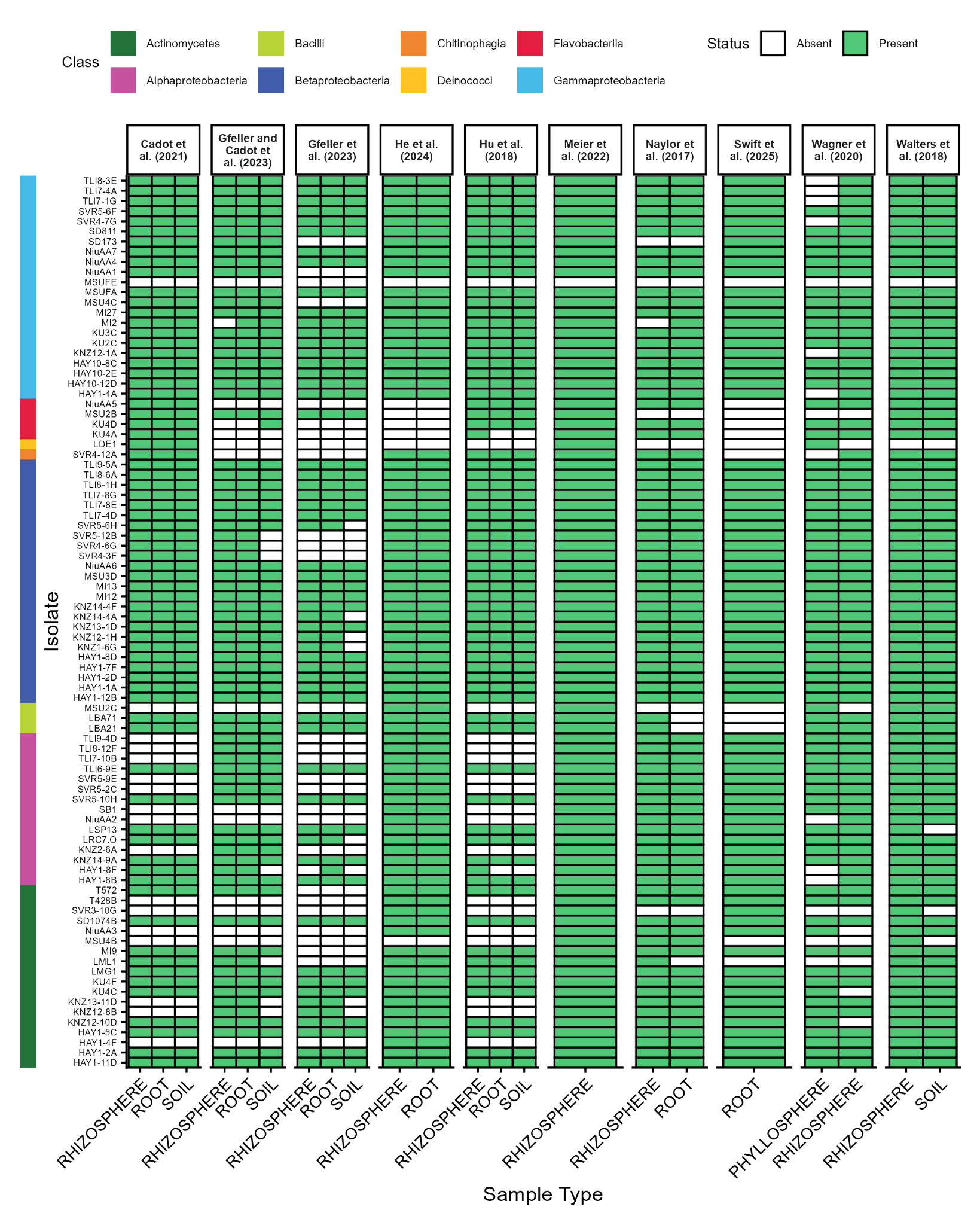
**Figure S5. The presence of ZeaMiC isolates in 16S rRNA gene maize microbiome datasets at 97% identity.** Significant alignments were filtered using a 97% identity threshold. For datasets containing multiple maize microbiome compartments, each compartment is shown as an individual subset within the study panel. ZeaMiC isolate labels are arranged by class (see Table S4 for full taxonomy of each ZeaMiC isolate).

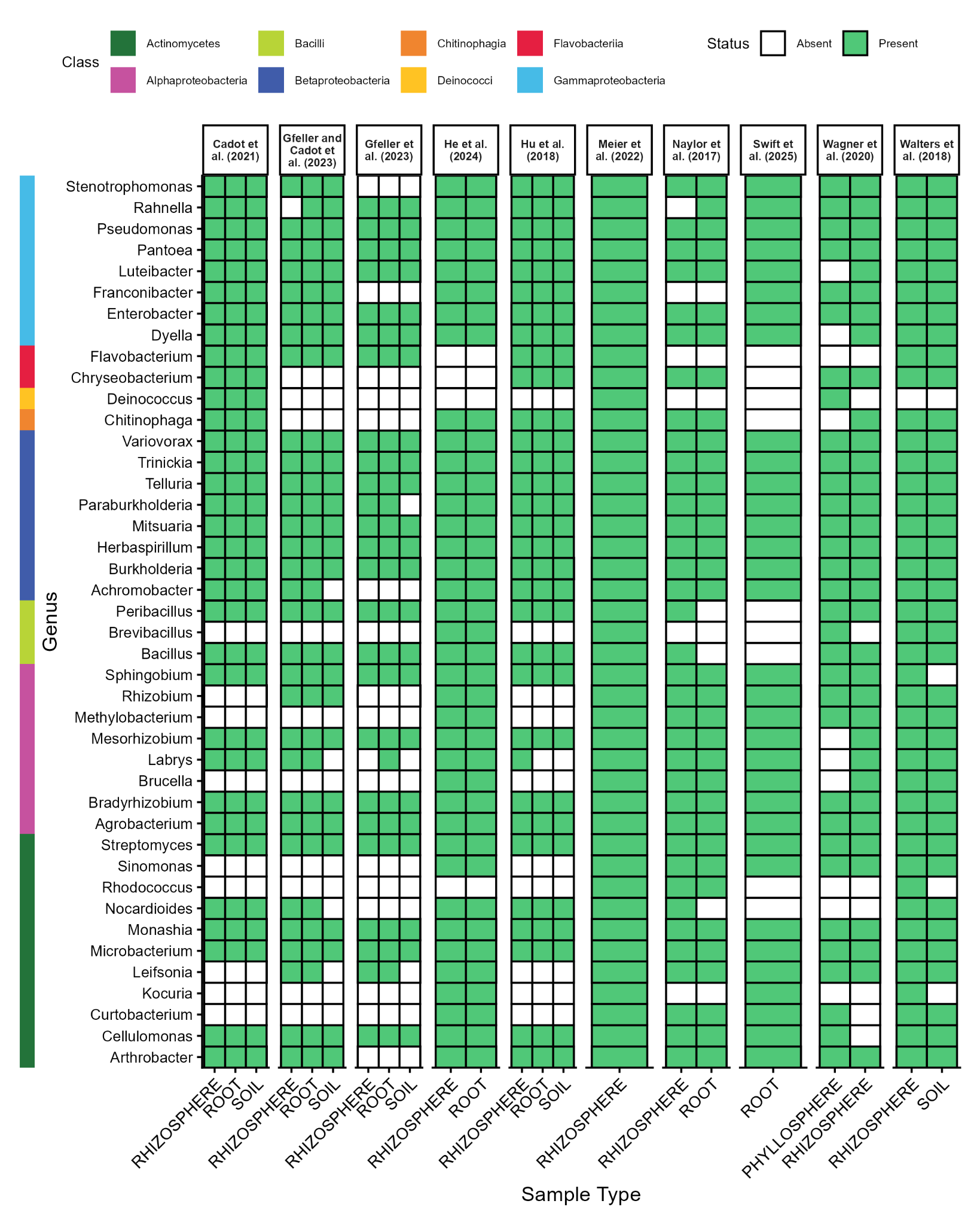

**Figure S6. The presence of ZeaMiC genera in maize 16S rRNA gene microbiome datasets at 97% identity.** Significant alignments were filtered using a 97% identity threshold. For datasets containing multiple maize microbiome compartments, each compartment is shown as an individual subset within the study panel. ZeaMiC genera labels are arranged by class (see Table S4 for full taxonomy of each ZeaMiC isolate).

### Supplementary Tables

**Table S1**. **Soils.** Soil sample sites information including the full site name, abbreviation, location (City, State, Country), sample date, and management type.

See supplemental table excel

**Table S2.** **Crude Collection.** Crude culture collection isolates information including originating soil source, method of isolation, and raw 16S rRNA gene sequences.

See supplemental table excel

**Table S3.** **Top 35 most abundant bacterial genera of Swift et al (2025).** List of the 35 most abundant bacterial genera observed in the root endosphere of 200 maize plants grown with Kansas prairie soil microbial inocula, as measured by 16S-V4 rRNA gene amplicon sequencing (Swift et al., 2025). Occupancy indicates the proportion of plants in which the genus was observed. Strain source indicates the location of the soil from which the strain was derived. HAY = Hays Prairie; KNZ = Konza Prairie; SVR = Smoky Valley Ranch; TLI = The Land Institute; KU = the University of Kansas Field Station; MSU = the Michigan State University Plant Pathology Research Farm; SBE = Sutton Bonington, England; TFL = Tarrafal, Cape Verde; SD = São Domingos, Cape Verde; SUI = Changins, Switzerland; AA = the Arnold Arboretum of Harvard University.

See supplemental table excel

**Table S4. Isolate Information.** List of ZeaMiC isolates with collection details (origin country/region, soil collection location, and soil type), associated maize genotype, isolate taxonomy and Gram stain, and public accession identifiers (NCBI BioProject, NCBI BioSample, NCBI genome accession when available, GenBank accession when available, and DSMZ accession number).

See supplemental table excel

**Table S5.** **Genome Annotation.** Microbial genes identified in each isolate genome from the curated list of plant-associated functions, genes, and KEGG Ortholog (KO) terms.

See supplemental table excel

**Table S6.** **DSMZ Bundles.** Isolates belonging to three cost-efficient bundles available through DSMZ.

See supplemental table excel

**Table S7.** **Robust detection of ZeaMiC taxa across maize microbiome studies.** For each dataset, we report the number of ZeaMiC isolates with significant ASV alignments and the relative abundance they contribute at the 97% and 99% sequence similarity thresholds. Contextual metadata are provided as well, including applied treatments, sampled compartments, microbiome source locations, and the number of samples and genotypes included. †Location refers to the cultivation site for field experiments, the soil source for container studies, or the origin of microbiome inocula. Presence and abundance values stratified by compartment are provided in Supplementary Figures S3-4 and Table S8.

See supplemental table excel

**Table S8.** **Compartment Coverage.** For each compartment profiled across the studies, we report the number of ZeaMiC isolates with significant ASV alignments and the relative abundance they contribute at 99% and 97% sequence similarity thresholds (shown above and below, respectively). Blank cells indicate that the compartment was not profiled as part of that study.

See supplemental table excel

**Table S9. ZeaMiC Comparison to Existing Datasets.** Common genera contributing the greatest relative abundance across datasets, defined as those present in at least 9 of 10 datasets. “Number of ASVs” denotes the number of individual ASVs recovered for a given genus, along with the studies from which each ASV originated. “Number of studies” refers to the number of unique studies in which an ASV was recovered. Mean occupancy and relative abundance were calculated by averaging the study-level values for each ASV within a genus across studies. All compartments were utilized in the calculator of mean occupancy and relative abundance to provide a more generalized value.

See supplemental table excel

### References

**Cadot S, Guan H, Bigalke M, Walser J-C, Jander G, Erb M, van der Heijden MGA, Schlaeppi K**. **2021**. Specific and conserved patterns of microbiota-structuring by maize benzoxazinoids in the field. *Microbiome* **9**: 103.

**Callahan BJ, McMurdie PJ, Rosen MJ, Han AW, Johnson AJA, Holmes SP**. **2016**. DADA2: High-resolution sample inference from Illumina amplicon data. *Nature Methods* **13**: 581–583.

**Camacho C, Coulouris G, Avagyan V, Ma N, Papadopoulos J, Bealer K, Madden TL**. **2009**. BLAST+: architecture and applications. *BMC Bioinformatics* **10**: 421.

**Gálvez-Merchán Á, Min KH (Joseph), Pachter L, Booeshaghi AS**. **2023**. Metadata retrieval from sequence databases with ffq. *Bioinformatics* **39**: btac667.

**Gfeller V, Cadot S, Waelchli J, Gulliver S, Terrettaz C, Thönen L, Mateo P, Robert CAM, Mascher F, Steinger T, *et al.* 2023a**. Soil chemical and microbial gradients determine accumulation of root-exuded secondary metabolites and plant–soil feedbacks in the field. *Journal of Sustainable Agriculture and Environment* **2**: 173–188.

**Gfeller V, Waelchli J, Pfister S, Deslandes-Hérold G, Mascher F, Glauser G, Aeby Y, Mestrot A, Robert CA, Schlaeppi K, *et al.* 2023b**. Plant secondary metabolite-dependent plant-soil feedbacks can improve crop yield in the field (S Rasmann, MC Schuman, B Delory, and I Kaplan, Eds.). *eLife* **12**: e84988.

**He X, Wang D, Jiang Y, Li M, Delgado-Baquerizo M, McLaughlin C, Marcon C, Guo L, Baer M, Moya YAT, *et al.* 2024**. Heritable microbiome variation is correlated with source environment in locally adapted maize varieties. *Nature Plants* **10**: 598–617.

**Hu L, Robert CAM, Cadot S, Zhang X, Ye M, Li B, Manzo D, Chervet N, Steinger T, van der Heijden MGA, *et al.* 2018**. Root exudate metabolites drive plant-soil feedbacks on growth and defense by shaping the rhizosphere microbiota. *Nature Communications* **9**: 2738.

**Martin M**. **2011**. Cutadapt removes adapter sequences from high-throughput sequencing reads. *EMBnet.journal* **17**: 10.

**McMurdie PJ, Holmes S**. **2013**. phyloseq: An R Package for Reproducible Interactive Analysis and Graphics of Microbiome Census Data. *PLoS ONE* **8**: e61217.

**Meier MA, Xu G, Lopez-Guerrero MG, Li G, Smith C, Sigmon B, Herr JR, Alfano JR, Ge Y, Schnable JC, *et al.* 2022**. Association analyses of host genetics, root-colonizing microbes, and plant phenotypes under different nitrogen conditions in maize. *eLife* **11**: 1–26.

**Naylor D, DeGraaf S, Purdom E, Coleman-Derr D**. **2017**. Drought and host selection influence bacterial community dynamics in the grass root microbiome. *The ISME Journal* **11**: 2691–2704.

**Swift JF, Kolp MR, Carmichael A, Ford NE, Hansen PM, Sikes BA, Kleiner M, Wagner MR**. **2025**. Drought stress homogenizes maize growth responses to diverse natural soil microbiomes. *Plant and Soil* **509**: 181–199.

**Wagner MR, Roberts JH, Balint‐Kurti P, Holland JB**. **2020**. Heterosis of leaf and rhizosphere microbiomes in field‐grown maize. *New Phytologist* **228**: 1055–1069.

**Walters WA, Jin Z, Youngblut N, Wallace JG, Sutter J, Zhang W, González-Peña A, Peiffer J, Koren O, Shi Q, *et al.* 2018**. Large-scale replicated field study of maize rhizosphere identifies heritable microbes. *Proceedings of the National Academy of Sciences* **115**: 7368–7373.

**Wang Q, Cole JR**. **2024**. Updated RDP taxonomy and RDP Classifier for more accurate taxonomic classification. *Microbiology Resource Announcements* **13**: e01063-23.

**Wang Q, Garrity GM, Tiedje JM, Cole JR**. **2007**. Naïve Bayesian Classifier for Rapid Assignment of rRNA Sequences into the New Bacterial Taxonomy. *Applied and Environmental Microbiology* **73**: 5261–5267.
